# Cumulative dose responses for adapting biological systems

**DOI:** 10.1101/2024.11.22.624851

**Authors:** Ankit Gupta, Eduardo D. Sontag

## Abstract

This paper introduces the notion of cumulative dose response (cDR). The cDR is the area under the plot of a response variable, an integral taken over a fixed time interval and seen as a function of an input parameter. This work was motivated by the accumulation of cytokines resulting from T cell stimulation, where a non-monotonic cDR has been observed experimentally. However, the notion is of general applicability. A surprising conclusion is that incoherent feedforward loops studied in the systems biology literature, though capable of non-monotonic dose responses, can be mathematically shown to always result in monotonic cDR.

## 1 Introduction

The capability to adapt and to formulate appropriate responses to environmental cues is a key factor for the survival of life, at every level from individual cells, to organisms, to societies [1, 2, 3, 4]. A delicate balance is needed in this process: organisms maintain tightly regulated levels of vital quantities, even in the face of variations to be counteracted upon, a property sometimes called homeostasis or adaptation, all the while being able to detect and react to changes in the environment. Underlying adaptation at the cellular level are dynamical signal transduction and gene regulatory networks that measure and process external and internal chemical, mechanical, and physical conditions (ligand, nutrient, oxygen concentrations, pressure, light, temperature) eventually leading to changes in metabolism, gene expression, cell division, motility, and other characteristics. These mechanisms enable organisms to display transient responses that gradually return to a baseline activity level when presented with relatively constant input stimulation, a phenomenon usually called “perfect” or “exact” adaptation [5].

### 1.1 Dose response and cumulative dose response

In this paper, we continue the study of adaptation mechanisms, with an emphasis on monotonicity properties of an output or reporter variable as a function of an input. We are concerned with ordinary differential equation systems that model the interactions of several species, and where there is an “input” (which might represent the dose of a drug or of a genetic inducer), whose level is quantified by the variable *u*, and an “output” that is time dependent, and whose magnitude at time *t* is represented by *y*(*t*). The input *u* will be assumed to be constant, and we write *y*_*u*_(*t*) to highlight the dependence of the output on both the input and time. Figure 1 shows three typical responses that one might observe experimentally (in the figure, we write *y*_*i*_(*t*) instead of 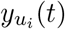, in order to simplify notations).

**Figure 1:**
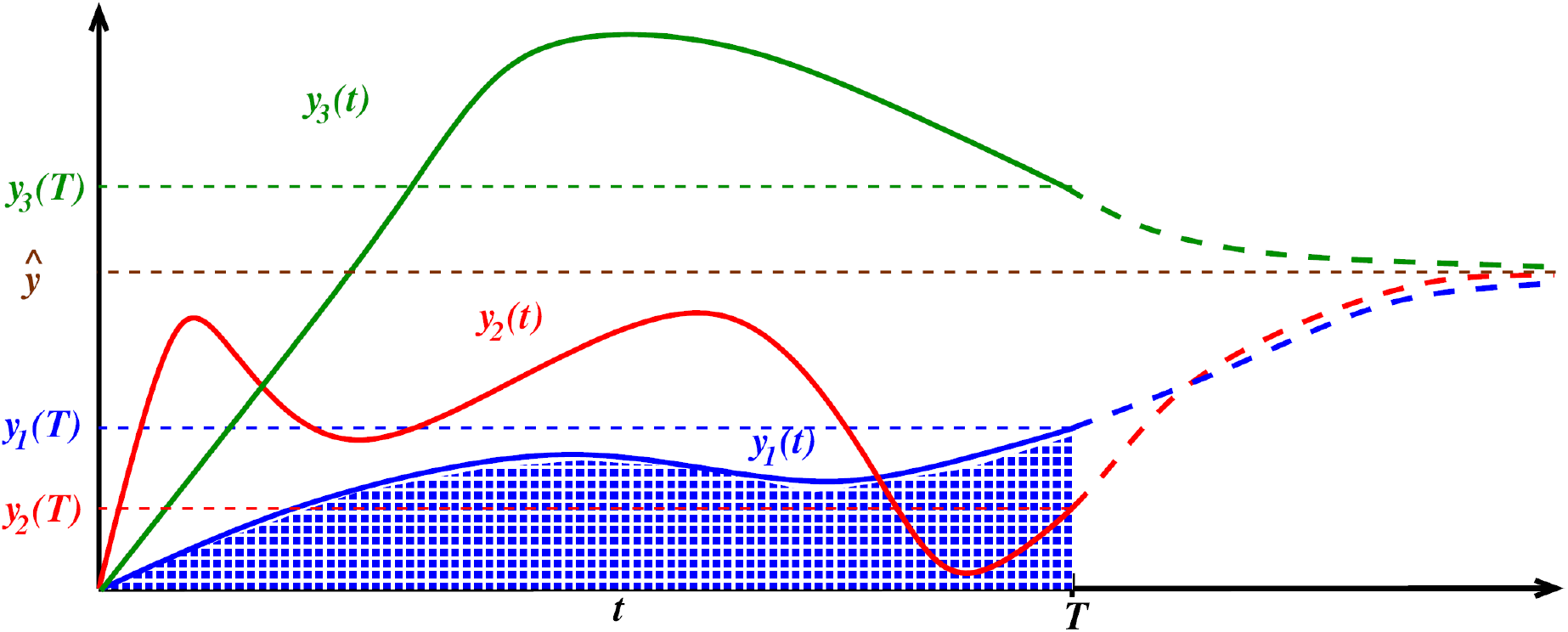
Response functions *y*(*t*) plotted against time. As an illustration, three outputs *y*_1_(*t*), *y*_2_(*t*), *y*_3_(*t*) are shown, corresponding to three inputs *u*_1_, *u*_2_, *u*_3_ respectively, and their values at a specified time *T* are shown on the vertical axis. In an adaptive system, the values of all the *y*_*i*_(*t*)’s converge to the same value *ŷ* when *t* → ∞. For each *y*_*i*_(*t*), solid curves are used for behaviors until the specified time *t* = *T*, and dashed lines for the continuation to *t* = ∞. The area of the shaded region represents the integral 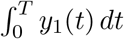.

The dependence on the initial state will not be indicated explicitly; the initial values of all the species will be fixed at values to be discussed.

One defines (“perfect”) adaptation to constant inputs as the property that, no matter what is the actual value of the input, the numerical value of *y*_*u*_(*t*) for large *t* is the same, that is, lim_*t*→∞_ *y*_*u*_(*t*) = *ŷ* for some fixed value *ŷ* which does not depend on the particular input *u* (see an illustration in Figure 1). This value represents a habituated or no-response state, as one achieves when presented with a constant background noise or level of light. In engineering terms, an adapting system is a “high pass filter” that essentially acts on a derivative of the input. Adaptation, by definition, is an *asymptotic* property, since it ignores finite-time behavior. On the other hand, *transient* behaviors, particularly how *y*_*u*_(*T*) varies with *u* at a fixed time *T*, are often of interest. (What is be the concentration of a drug in a tumor microenvironment, after 1hr, as a function of the drug dose? What is the size of a tumor after 60 days of the start of therapy, as function of the drug dose? What is the size of the pool of infected individuals in an epidemic population model, as a function of transmission parameters?)

We will call *y*_*u*_(*T*), viewed as a function of *u* the *dose response* of the given system, and denote it as DR(*u, T*). One may perform experiments, exciting a system with different input values *u*, and measure *y*_*u*_(*T*) as the final output value, thus obtaining a plot of *y*(*T*) versus *u*. The left panel of Figure 2 shows several dose responses obtained from time-resolved data such as presented in Figure 1.

**Figure 2:**
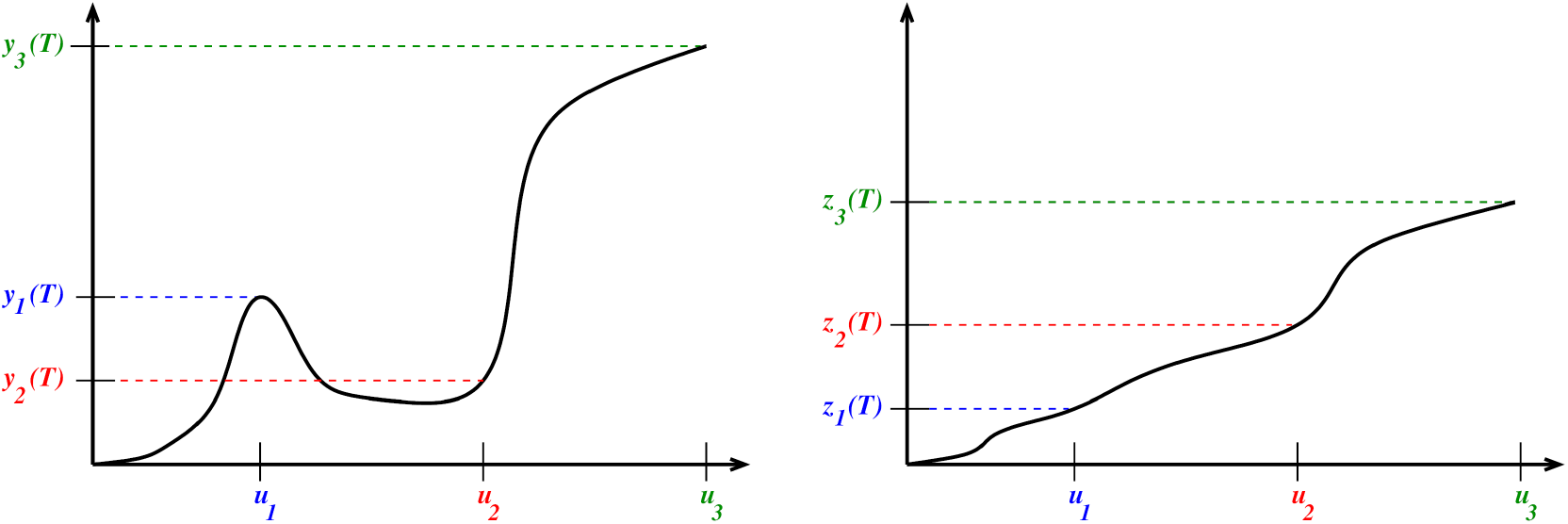
Left: Dose response (DR) at time *T*, obtained from time-resolved data as in Figure 1. Again, the vertical axis represents the evaluation of responses *y*(*t*) at a specific time *t* = *T*, but now these values are plotted against the respective input *u*, instead of against time. In this instance, DR is not a monotonic function; for example, *u*_1_ *< u*_2_ but *y*_1_(*T*) *> y*_2_(*T*). Right: Cumulative dose response (cDR) at time *T*; now the vertical axis shows the integral (area under the curve) 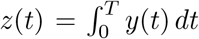 of the response, again plotted against the input *u*. For example, *z*_1_(*t*) is the area shaded in blue in Figure 1. This particular cDR is monotonic. For example, *u*_1_ *< u*_2_ *< u*_3_ and *z*_1_(*T*) *< z*_2_(*T*) *< z*_3_(*t*), because in Figure 1 the area under the red curve *y*_2_(*t*) is larger than the area under the blue curve *y*_1_(*t*), even though the final value *y*_2_(*T*) is smaller than *y*_1_(*T*).

If the turnover of *y*(*t*) is slow, the molecules or other objects represented by *y*_*u*_(*t*) may accumulate, for instance, in a particular tissue or the bloodstream. It is often the case that one can only measure experimentally, and that a phenotypical response only depends on, the accumulated value or integral 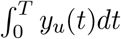, which we call the *cumulative dose response* and denote by cDR(*u, T*). The right panel of Figure 2 shows several cumulative dose responses obtained from data as in Figure 1. Indeed, this paper was motivated by our previous research in immunology that measured cDR’s. We discuss that motivation next.

### 1.2 Motivation: T cell recognition

Adaptation is central to immunity. In particular, T cells must react to stimulation by pathogens and cancers, yet limit their response in order to maintain self-tolerance and avoid autoimmune reactions. T cell activation is triggered by the binding to T cell receptors (TCRs) to peptide major-histocompatibility complex (pMHC) antigens. Activation results in the production of signaling molecules (cytokines) which in turn may recruit other immune components.

The study in [6] examined the response of immune CD8+ T cells to external antigen inputs, demonstrating perfect adaptation across a wide range of antigen affinities. Specifically, in the experiments in [6] involved stimulating primary human CD8+ T cells (with the c58c61 TCR) with recombinant pMHC antigen 9V immobilized on plates, which served as the input *u* at various constant concentrations. This antigen is a cancer peptide routinely used in studies of T-cell binding and antigen discrimination. The cumulative amount of cytokine TNF-*α* (the output *y*(*t*)) secreted into the culture medium was measured.

Figure 3 shows the cDR when one averages the results of three biological replicates. It shows the average cumulative TNF-*α* abundance 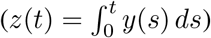 plotted against several constant concentrations *u*, measured at various times (*t* = 1 to 8 hours). Note the non-monotonic, and even somewhat oscillatory, behavior.

**Figure 3:**
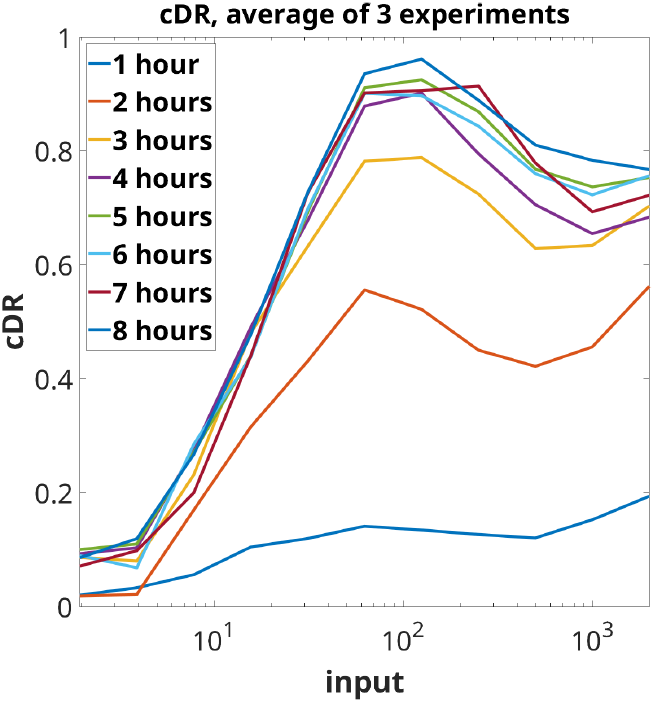
Cumulative dose response based on average of three experiments. Plot uses experimental data from [6] (see also top left panel of Figure 1 in that paper). Horizontal axis denotes concentrations of the input (in units of ligand in ng/well).

The work in [6] thus raises the question of what network motifs are capable, at least for suitable parameters, to exhibit perfect adaptation as well as non-monotonic cDR as seen in these experiments. It was speculated, on the basis of numerical exploration, that incoherent feedforward loops cannot result in non-monotonic cDR and thus cannot explain T cell adaptation as measured by accumulated cytokines, unless a thresholding mechanism is imposed.

Our main results in this paper confirm in a mathematically rigorous way that, indeed, the main two common types feedforward loops (called “IFFL1” and “IFFL2” below) can *never* exhibit such behaviors, because their cDR’s are *always monotonic*. This is especially surprising for one of them (“IFFL1”) because for such systems the DR itself can be non-monotonic, yet the cDR is monotonic, as in the cartoon illustrations in Figure 2. To see this with an example, consider the following system of two differential equations:

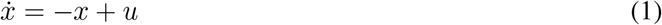

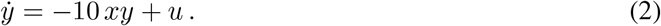

This is an adapting system: for any given constant input *u* > 0, the steady states are 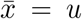 and 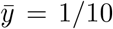, which is independent of *u*, so that the steady state output *ŷ* = 1*/*10 is independent of *u*. Figure 4 shows plots of the DR (non-monotonic) and the cDR (monotonic).

**Figure 4:**
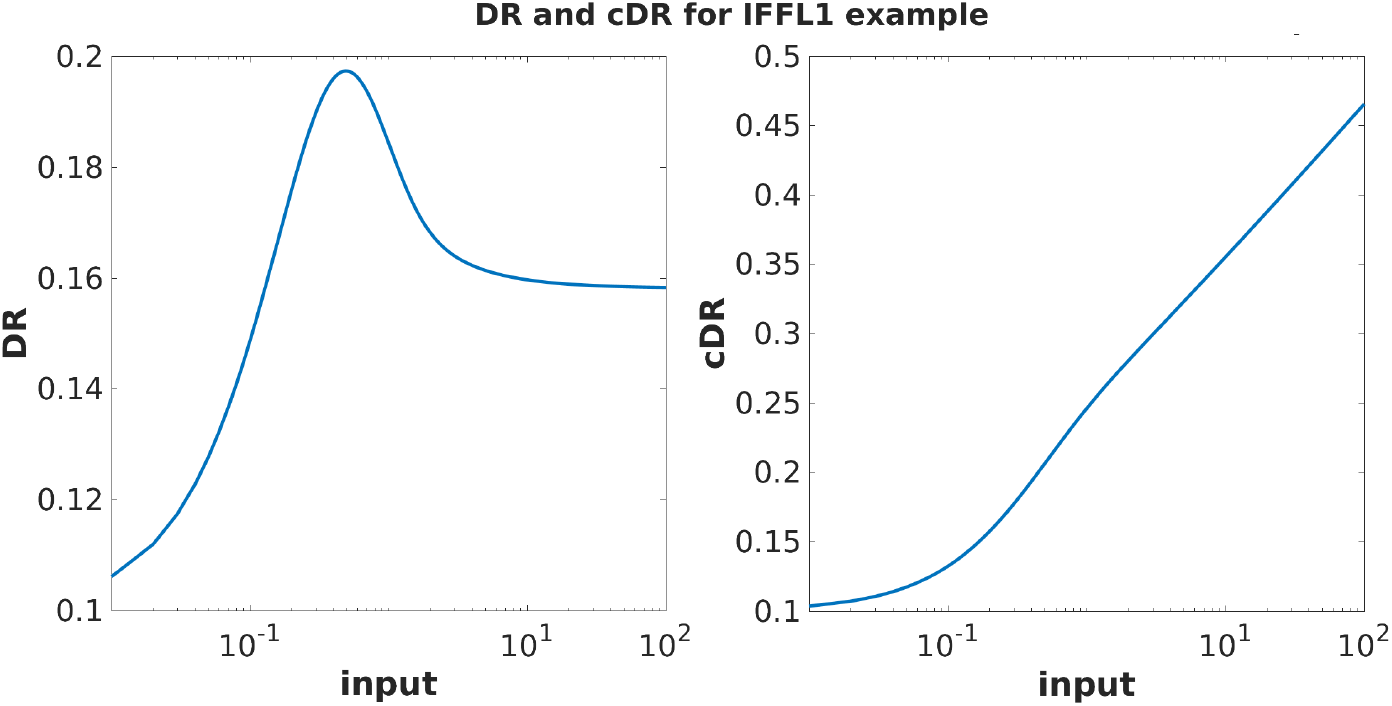
Plots of DR (*y*(*t*)) and cDR 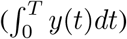 for the example in Equations (1-2). The initial conditions are *x*(0) = 0, *y*(0) = 1*/*10, and the time horizon is *T* = 1.5. Using logarithmic scale on inputs, for comparison with experimental plots. Observe that the DR is non-monotonic, yet, surprisingly, the cDR is monotonic. Our main theorem proves monotonicity of the cDR in general, for all IFFL1 and IFFL2 systems.

We complement the result for feedforward loops with the new finding that, on the other hand, the standard *nonlinear integral feedback* for adaptation (“IFB” below) is indeed capable of showing non-monotonic cDR, and thus is potentially a mechanism that is consistent with the experimentally observed non-monotonic cDR. To see this with an example, consider the following system of two differential equations:

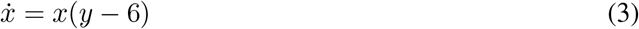

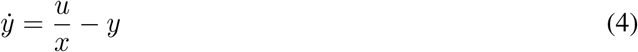

This is also an adapting system: for any given constant input *u >* 0, the steady states are 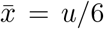 and 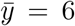, so that the steady state output *ŷ* = 6 is independent of *u*. Figure 5 shows plots of DR (non-monotonic) and cDR (monotonic).

**Figure 5:**
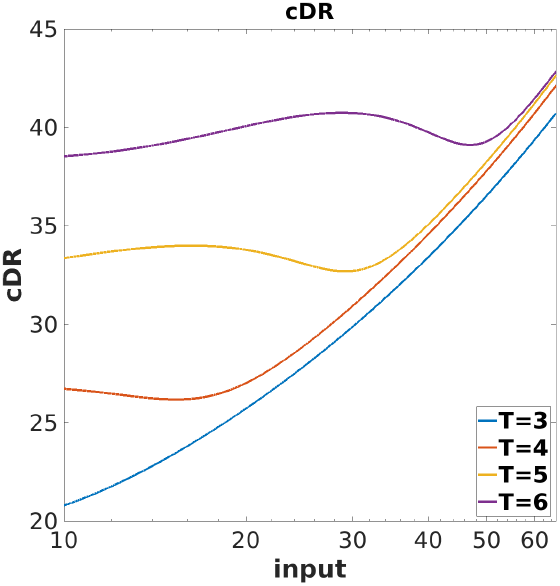
Plot of cDR 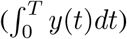 for the integral feedback example in Equations (3-4). The initial conditions are *x*(0) = 0.1, *y*(0) = 6, and time horizons shown are *T* = 3, 4, 5, 6. Observe that, just as with the experimental data, the cDR is more monotonic (on the shown ranges, at least) for smaller time horizons *T*. Using logarithmic scale on inputs, for comparison with experimental plots.

Back to the experimental data, one may ask whether the T cell experiments point to adaptation, that is, if *y*_*u*_(*t*) is independent of *u* for large *t*. Since only 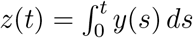 is experimentally available, we need to estimate the output *y*(*t*) by taking time-derivatives of *z*(*t*). To obtain a more meaningful estimate than would be obtained from the averages shown in Figure 3, we consider instead the separate plots of *z*(*t*) from each experiment. The top panel in Figure 6 shows again the cumulative dose responses for various times (*t* = 1 to 8 hours), but now with separate plots from each experiment, starting from the data that was used to generate Figure 1 in the SI of [6].

**Figure 6:**
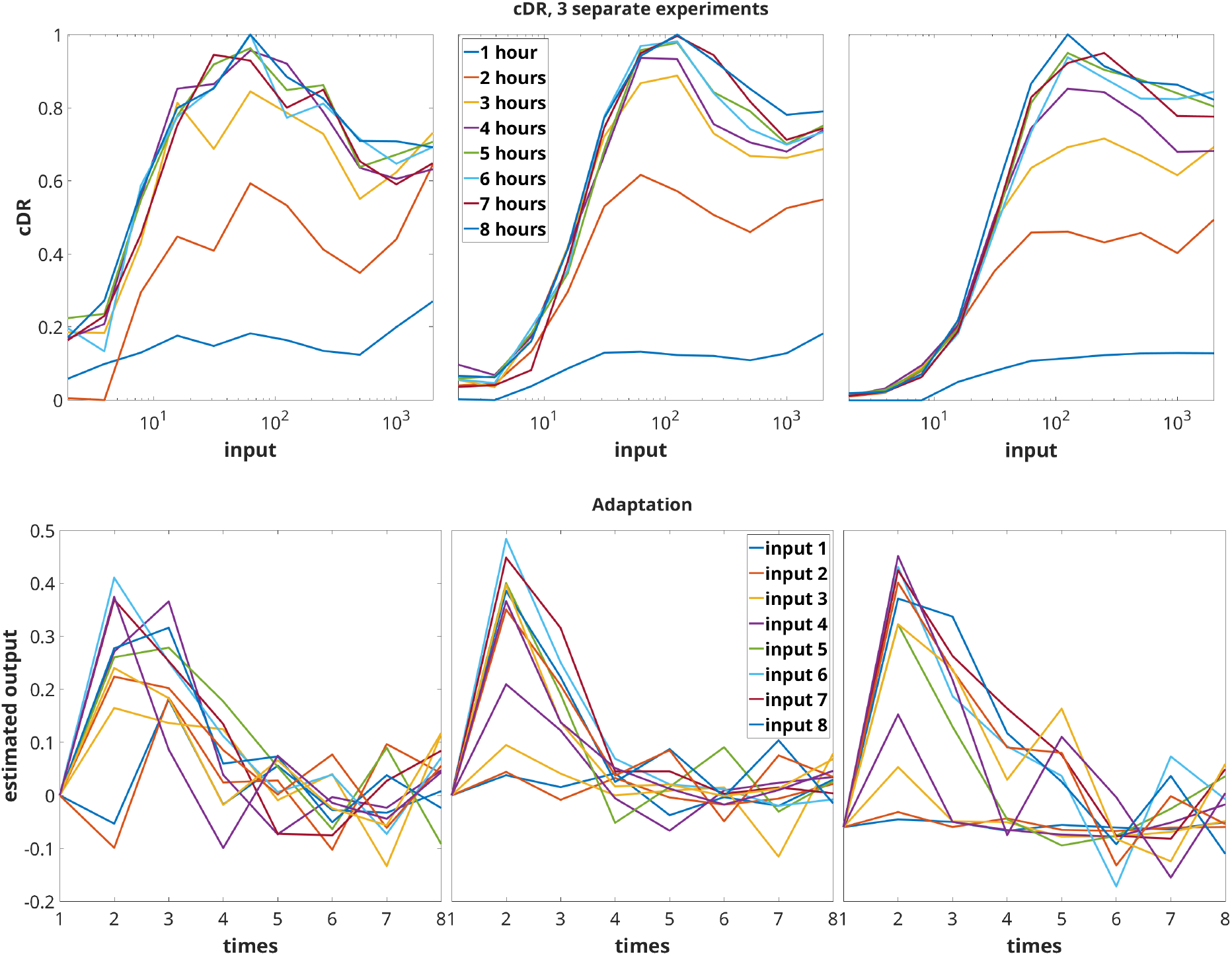
Top: cDR plots of individual experiments, and measured at different times. Bottom: Adaptation behavior in individual experiments. Output *y*(*t*) is estimated from individual cumulative *z*(*t*) plots in respective top panels.

Using first-order differences, and imposing a zero value at the start of the experiment: *y*(0) = 0, we then derived estimates of *y*(*t*) for the various input values and the three experimental replicates shown in Figure 6. See the bottom panel of Figure 6. These estimates are very rough, because the time steps are large (1 hour), and in any event experimental data is subject to noise. Nonetheless, this data strongly suggests that adaptation, in the sense that the output *y*(*t*) approaches as time increases a value (in this case *y* = 0) which is the same no matter what the input (drug dose). The estimated negative values of *y*(*t*) at certain time points are likely due to numerical errors or to the fact that there is some cytokine present in the experimental wells which does not arise from the stimulation, so that the values shown are relative to this baseline. In addition to adaptation, the data strongly suggests that the system, or at least its integrated output, is *scale-invariant* (performs “fold change detection”) in the sense of [7, 8], at last for large enough input values: the transient outputs are roughly the same (for inputs 3-8), which is a property of systems IFFL2 and the integral feedback system as discussed below.

### 1.3 Review: network motifs for adaptation

As discussed, (“perfect”) adaptation to constant inputs means that, no matter the value of an input, the value of *y*_*u*_(*t*) for large *t* will be the same. Generally speaking, adaptation requires one of two mechanisms for adaptation: incoherent feedforward or negative feedback [9, 10, 11]. While the concepts we introduce are broadly applicable, our examples will focus on systems with only two variables for clarity.

*Incoherent feedforward loops (IFFL)* are a type of network motifs that are capable of adaptation [12, 13]. In an IFFL, the input *u* induces formation of the reporter *y* but also acts as a delayed inhibitor, through one or more intermediary control variables. Feedforward motifs are statistically overrepresented in biological systems from bacterial to mammalian cells [14, 15]. IFFL’s have been argued to underlie mechanisms involved in such varied contexts as microRNA-mediated loops [16], MAPK pathways [17, 18], insulin release [19, 20], intracellular calcium release [21, 22], *Dictyostelium* and neutrophil chemotaxis [23, 24], NF-*κ*B activation [25], and microRNA regulation [26], as well as metabolic regulation of bacterial carbohydrate uptake and other substrates [27, 28]. IFFL’s may also play a role in immunology, enabling the recognition of dynamic changes in antigen presentation [29], and have been employed in synthetic biology in order to control protein expression under DNA copy variability [30, 31]. The paper [32] provided a large number of additional references, and carried out a computational exploration of IFFLs that lead to non-monotonic dose responses and/or adaptation.

In *integral feedback loops (IFB)*, the intermediate variable or variables provide a type of memory that integrates the “error” between *y*(*t*) and a steady-state value *y*_0_. IFB’s arise in biological systems ranging from *E. coli* chemotaxis [33] and regulation of tryptophan [34] to human physiological control such as blood calcium homeostasis [35] or neuronal control of the prefrontal cortex [36] to synthetic circuits for adaptation [37, 38, 39]. We remark that integral feedback is in a sense universal for adaptation, because nonlinear changes of coordinates can recast IFFLs that adapt into integral feedback form [40], but these coordinate changes may have no physical interpretation and hence lack interpretability. Moreover, IFBs are known to provide extra degrees of robustness to the adaptation property because, unlike the IFFL, the underlying mechanism can sense and correct for perturbations to the output variable *y*(*t*). This theme is explored in greater detail in recent papers, where it is shown that IFBs arise naturally when one considers adapting circuits that exhibit a *maximal* form of robustness [41] or robustness that is independent of the reaction kinetics [42]. The results in these papers apply to arbitrarily-sized networks, and they explicitly identify the network species that create the integral feedback required for adaptation.

### 1.4 Three paradigmatic examples of adapting systems

In this paper, we will consider three types of two-species systems, shown schematically in Figure 7. These are the systems studied in [7, 8]. In all these examples, *u* refers to a positive constant input, *x* is the concentration of a “controller” species, and the output variable *y* is the concentration of the “regulated” or output species.

**Figure 7:**
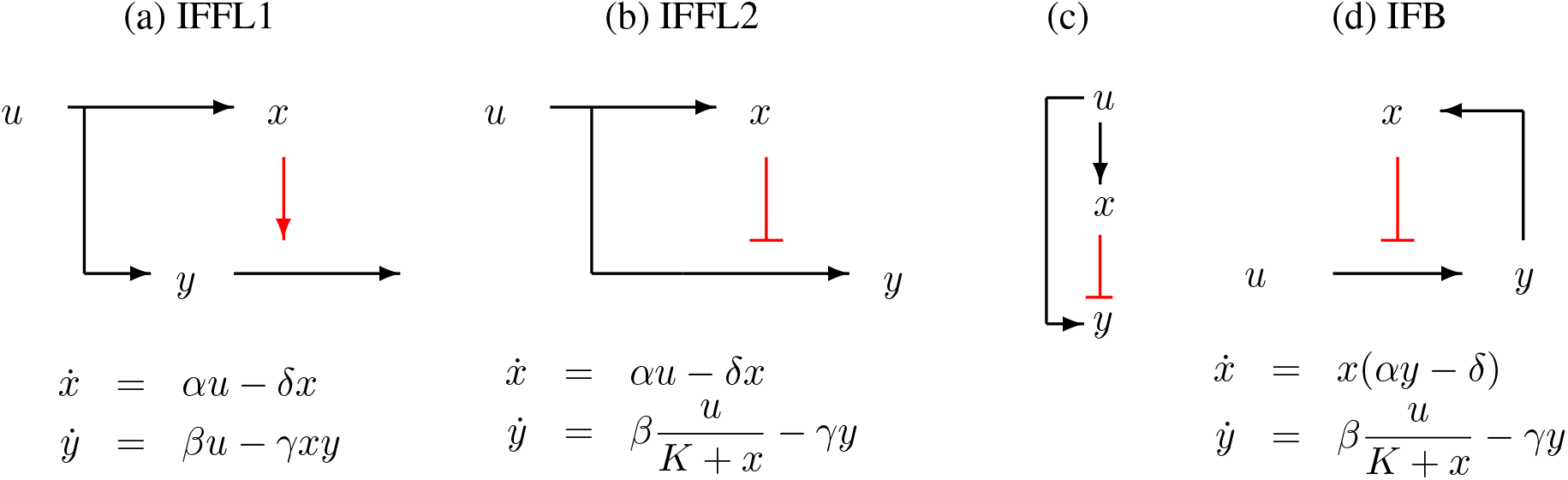
Three examples of systems: (a) is an IFFL with degradation enhancement, (b) is an IFFL with production inhibition, and (d) is an integral feedback system. The input *u* is assumed to be a positive constant, and *x, y* are abundances of a quantity of interest such as concentration of a protein or mRNA. Arrows “ → “ indicate positive effect (activation) and blunt edges “ ⊣ “ denote negative effects (inhibition). Both (a) and (b) have the same qualitative relation, schematically represented by diagram in (c), between activation of *x* and *y* by the input *u*, and inhibition of *y* by *x*, but their dynamical properties are very different. Dynamics can be described by pairs of differential equations for the abundances *x*(*t*) and *y*(*t*) as a function of time, as shown below the diagrams. State variables *x*(*t*) and *y*(*t*) are taken to be nonnegative. When *K* = 0, it is assumed that *x*(*t*) *>* 0 for all *t*. In these equations, *α, β, γ, δ*, and *K* are all positive constants.

### 1.5 Steady states and perfect adaptation

Let us compute the steady states, obtained by setting the right-hand sides of the differential equations to zero, for the systems shown in Figure 7. For the IFFL1 system (a) we have:

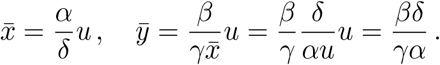

For the IFFL2 system (b) we have:

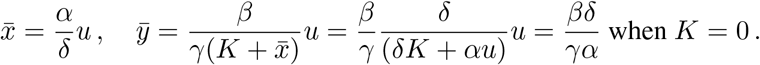

Finally, for the IFB system (c) we have that there are two types of steady states:

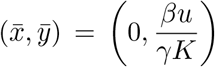

and, for nonzero 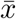 and assuming 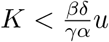 (for example, if *K* = 0):

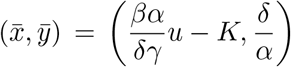

In all three cases, if *K* = 0, the system is perfectly adapting.

### 1.6 Scale-invariance

In addition, when *K* = 0 both systems IFFL2 and IFB have the scale-invariance (or “fold-change detection”, FCD) property [7, 8]. This means that the output variable *y*(*t*) satisfies the same differential equation, independently of rescalings of the input *u* by any constant factor *p*, as shown by the following simple calculation:

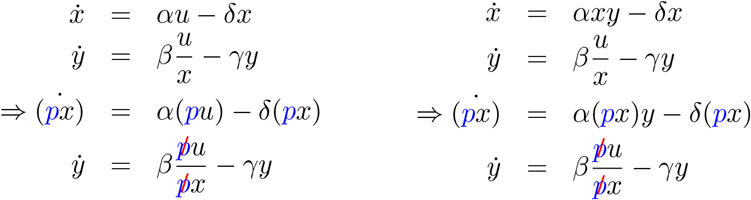

System IFFL1, in contrast, admits no such symmetries.

### 1.7 Stability

For both systems IFFL1 (a) and IFFL2 (b) in Figure 7, the respective steady states 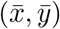 are globally asymptotically stable with respect to initial conditions in the positive quadrant *x >* 0, *y >* 0. This is very simple to show. The variable *x*(*t*) is the solution of a one-dimensional stable linear system, hence converges exponentially to 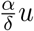. The variable *y* is the solution of a time-dependent linear system, with a constant negative rate −*γ* for IFFL2, and a rate for IFFL1 which converges to the strictly negative number 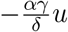, and hence also exponentially converges to its steady state value. (See [13] for details, as well as similar results for other IFFL configurations.)

The proof of stability for the feedback system IFB (d) in Figure 7 requires more work. We proceed by extending the proof from [8], which covered only the case *K* = 0. We will assume that 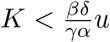, which holds in particular if *K* = 0. We want to global asymptotic stability with respect to initial conditions with *x*(0) positive and *y*(0) non-negative (or even arbitrary), when *u* is a positive constant, for the two-dimensional system IFB evolving on ℝ_*>*0_ *×* ℝ with equations 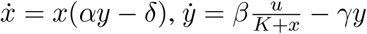. It is convenient to change coordinates 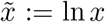 and 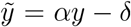, so that we reduce to the study of the system in ℝ^2^ with equations (dropping tildes):

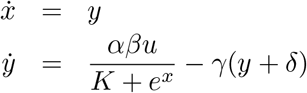

and we wish to prove the global asymptotic stability of the unique steady state

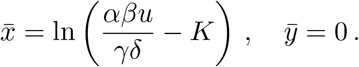

Introducing 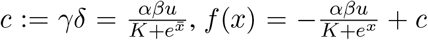, and 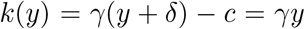, we can write our system as

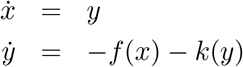

with 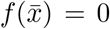 and 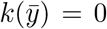. In other words, we have a mass-spring system 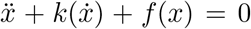 with nonlinear damping 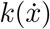 and nonlinear spring constant *f* (*x*). This suggests the use of energy as a Lyapunov function. The map *f* is strictly increasing, positive for 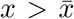 and negative otherwise, and similarly for *k* with respect to 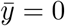. Let us define

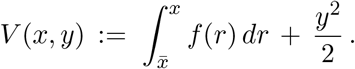

By definition, 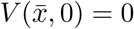 and *V* (*x, y*) *>* 0 for all 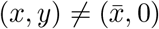. We also have that 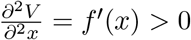, and 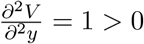, (and mixed partial derivatives are zero), *V* is a strictly convex, and thus a proper (also called radially unbounded or coercive) function, thus a Lyapunov function candidate. The derivative of *V* along trajectories is

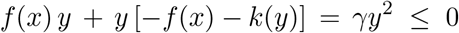

for all (*x, y*), and if this derivative only vanishes identically along a trajectory, then *y*(*t*) ≡ 0, which in turn implies, when substituted into 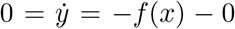 that *f* (*x*(*t*)) ≡ 0, i.e. that 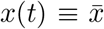. The LaSalle Invariance Principle (see e.g. [43]) then allows us to conclude global asymptotic stability.

### 1.8 Outline of paper

Denoting the input-dependent dynamics as (*x*_*u*_(*t*), *y*_*u*_(*t*)) for *t* ≥ 0, we define the **dose response (DR)** and the **cumulative dose response (cDR)** at time *T* as

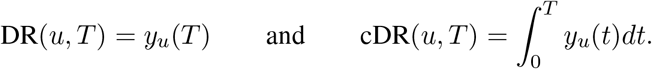

In each example, our aim is to determine whether the mapping *u* ↦ cDR(*u, T*) is monotonically increasing. If this monotonicity does not hold universally, we seek to identify sufficient conditions under which it does. It is straightforward to note that if the map *u* ↦ DR(*u, T*) is monotonically increasing, then the same holds for the map *u* ↦ cDR(*u*, ↦ *T*). Observe that if the system is linear, both DR(*u, T*) and cDR(*u, T*) will be linear (and therefore monotonic) functions of u. Hence, non-monotonicity is exclusively a property of nonlinear systems.

Let us review the specific examples that we consider. The first is the IFFL1 system shown in Figure 7(a) with equations given by

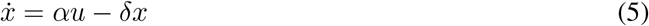

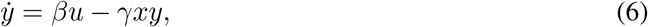

with initial state *x*(0) = *x*_0_ and 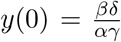. The initial state for *x* is arbitrary nonnegative, while the initial state for *y* is its steady-state value which is independent of *u >* 0.

The second example is the IFFL2 system (see Figure 7(b)) given by

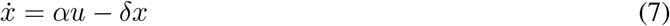

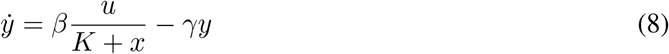

where the initial states *x*_*u*_(0) = *x*_0_ *>* 0 and *y*_*u*_(0) = *y*_0_ *>* 0 are arbitrary.

As the final example we have the IFB system (see Figure 7(c)) given by

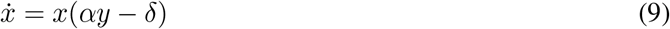

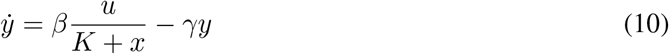

with initial states *x*(0) = *x*_0_ *>* 0 (arbitrary) and 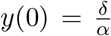, which is the steady-state for *y* which is independent of *u*.

We remark that for all of these examples, *u* ↦ DR(*u, T*) (and therefore also *u* ↦ cDR(*u, T*)) is monotonically increasing for *small T*. This is because 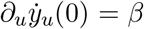 for IFFL1, and 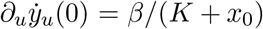 for IFFL2 and IFB. From 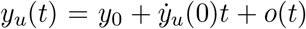, it follows that ∂_*u*_*y*_*u*_(*T*) ≈ *βT* and ∂_*u*_*y*_*u*_(*T*) ≈ (*βT*)*/*(*K* + *x*_0_) respectively, and both are positive. The situation is far less trivial for larger times *T*. The rest of this paper is organized as follows. In Section 2 we prove that for IFFL1, even though the map *u* ↦ DR(*u, T*) may not be monotonic, the map *u* ↦ cDR(*u, T*) is always monotonically increasing, irrespective of the values of *T*, *x*_0_, *δ* and *γ*. In Section 3 we show that the situation is much simpler for IFFL2, in the sense that both DR(*u, T*) and cDR(*u, T*) are monotonically increasing functions of *u*, regardless of the choice of *T*, initial states and the model parameters. Lastly, in Section 4 we show that for the IFB system the map *u* ↦ cDR(*u, T*) is not monotonic in general and find a sufficient condition under which monotonicity holds.

## 2 Unconditional monotonicity of the cDR for IFFL1

We shall prove the monotonicity of map *u* ↦ cDR(*u, T*) in multiple steps.

### 1. Simplifying the system

As the first step, we simplify the system (5)-(6) to make it more amenable to analysis. Let (*x*_*u*_(*t*), *y*_*u*_(*t*)) be the solution of this system for *t* ≥ 0. We scale time by *δ*^−1^ and the state values by a suitable ratio to define

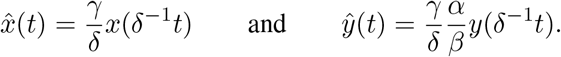

Then the dynamics of this rescaled system are given by

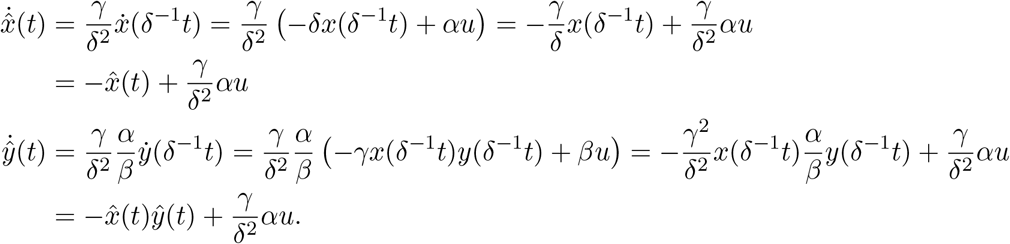

Therefore if we define

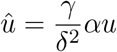

then 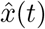, *ŷ*(*t*)) satisfies the ODEs

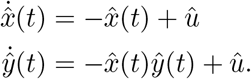

with initial states 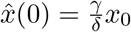 and *ŷ*(0) = 1. Let 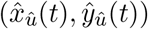, *ŷ*_*û*_(*t*)) be the solution of this system. To prove the result it suffices to show that the map

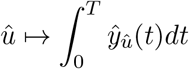

is monotonically increasing for any *T >* 0.

Henceforth we shall drop the hats for notational convenience and suppose that the dynamics is given by

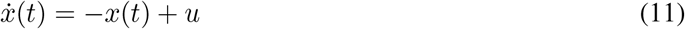

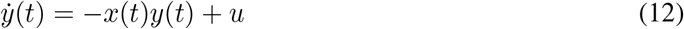

with initial state *x*(0) = *x*_0_ (arbitrary) and *y*(0) = 1. Letting (*x*_*u*_(*t*), *y*_*u*_(*t*)) be the solution of this system, we shall show that the cDR map

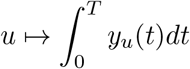

is monotonically increasing for any *T >* 0. In order to prove this we will prove that for any *T* and *u*

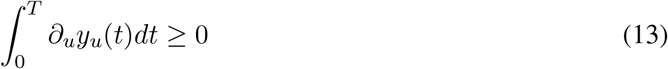

where ∂_*u*_ denotes the partial derivative with respect to *u*.

### 2. Deriving explicit expressions

As the second step, we develop explicit expressions for *x*_*u*_(*t*), *y*_*u*_(*t*), 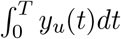, and 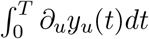. It is easy to see that

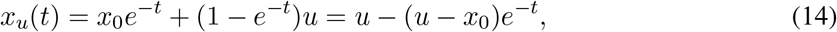

which also implies that

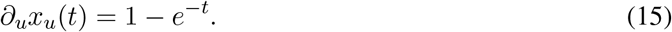

Note that ODE (12) can be written as

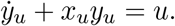

Multiplying with the integrating factor 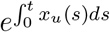 on both sides we obtain

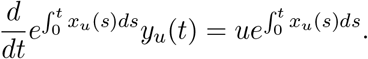

Integrating both sides and using *y*_*u*_(0) = 1 we get the usual variation of parameters formula:

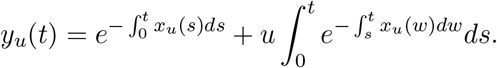

From (11) we know that

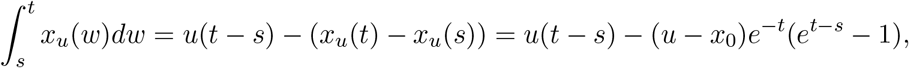

where the second relation follows from (14). Plugging this in the previous expression for *y*_*u*_(*t*) we get

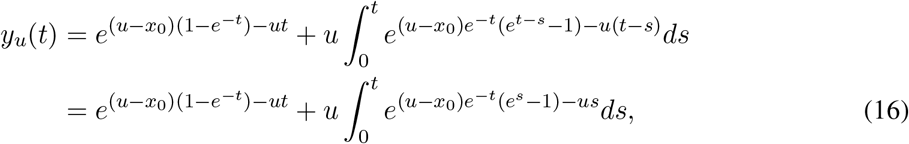

where to derive the last expression we have made a change of variable from (*t* − *s*) to *s*.

Note that by changing the order to integration in the second term in the r.h.s. below we obtain

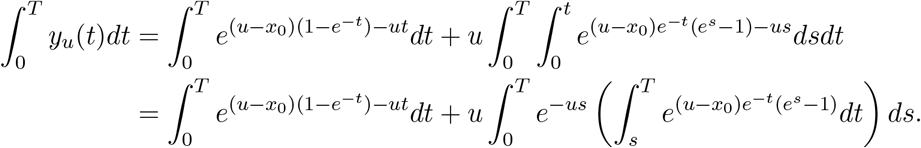

Making the change of variable *t* ↦ (*u* − *x*_0_)*e*^−*t*^(*e*^*s*^ − 1) we see that

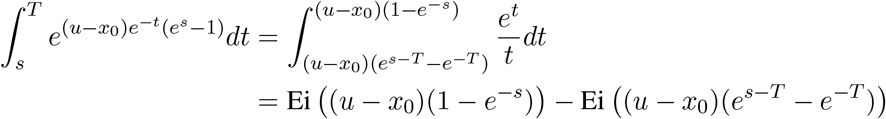

where Ei(*t*) is the special *Exponential Integral* function defined as

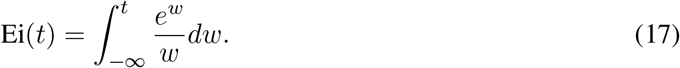

Plugging this in we obtain

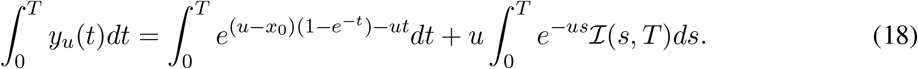

where

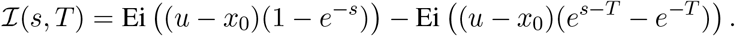

Observe that at *s* → *T*^−^ we have ℐ(*s, T*) → ℐ(*T, T*) = 0 and we claim that as *s* → 0^+^, we have ℐ(*s, T*) → *T*. To see this note that as *s* → 0^+^, both (1 − *e*^−*s*^) and (*e*^*s*−*T*^ − *e*^−*T*^) approach 0 and close to 0 we have

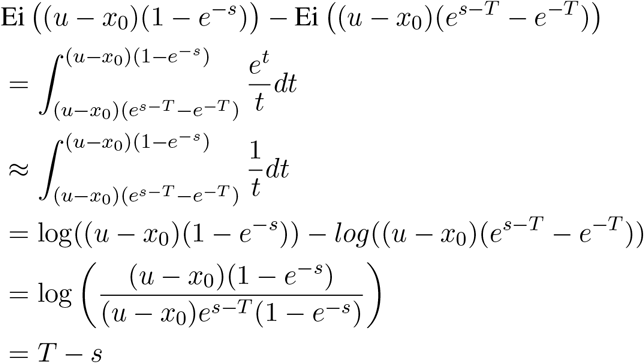

which is just *T* as *s* → 0+. By the definition of the Exponential Integral function (17)

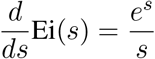

and hence by the chain rule

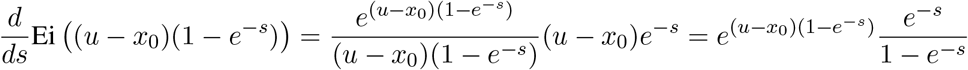

and

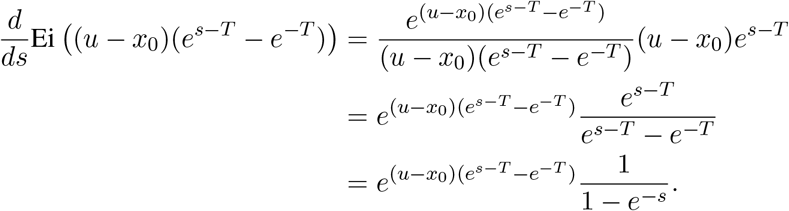

Using these expressions we can write the derivative of ℐ(*s, T*) as

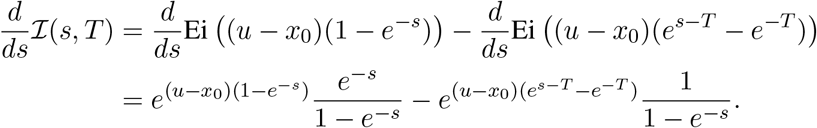

Therefore applying integration by parts to the second term in the r.h.s of (18) we get

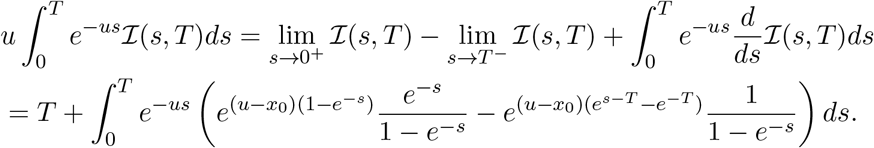

Upon substituting this term in the r.h.s. of (18) we obtain

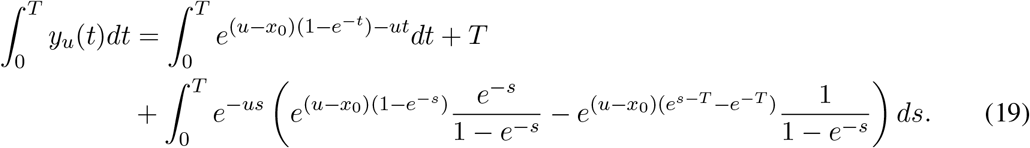

Note that by rearranging and simplifying, (19) can be expressed as

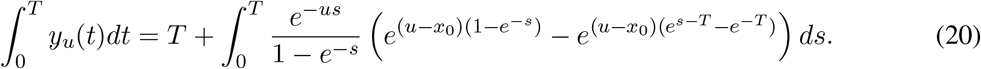

Differentiating (20) with respect to *u* we obtain

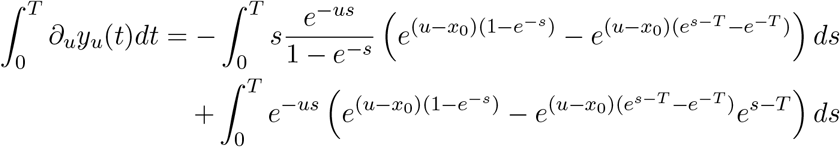

which can be rewritten as

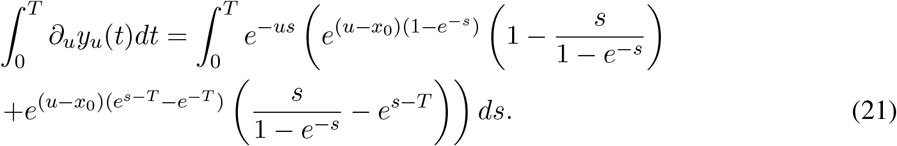

### 3. Two useful results

We now establish a couple of useful results that will help us in proving the monotonicity property of the cDR.

#### Lemma 1.

*Fix any positive T and κ, and for s* ∈ [0, *T*] *define functions*

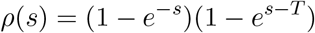

*and*

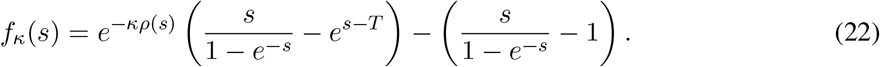

*Then the function f*_*κ*_(*s*) *can cross the x-axis only once in the interval* [0, *T*].

*Proof*. Note that *ρ*(*s*) = (1 − *e*^−*s*^)(1 − *e*^*s*−*T*^) = 1 + *e*^−*T*^ − *e*^−*s*^ − *e*^*s*−*T*^. Setting *f*_*κ*_(*s*) = 0 we get

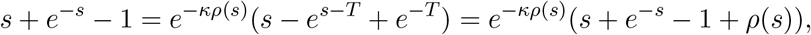

which upon re-arranging yields

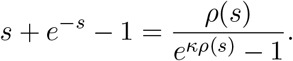

Let *g*_*κ*_(*s*) be the function

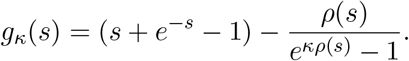

To prove the lemma it suffices to show that the function *g*_*κ*_(*s*) is monotonically increasing, which we shall show by proving that 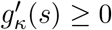 for all *s* ∈ [0, *T*]. For convenience let us define a function

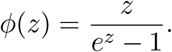

Then we can write *g*_*κ*_(*s*) as

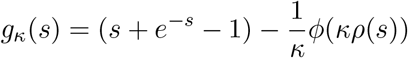

which shows that

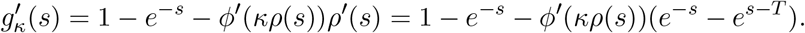

Suppose we can show that

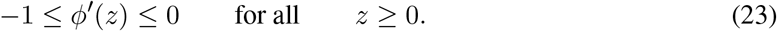

Then for 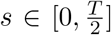 we have (*e*^−*s*^ − *e*^*s*−*T*^) *>* 0 and so 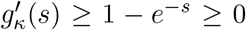. On the other hand for 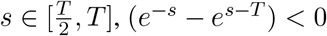 and using 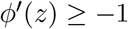, we obtain

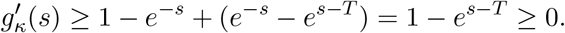

Hence to prove the lemma we just need to prove the inequality (23) to which we come to now. Note that

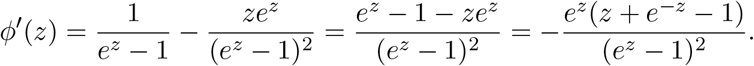

Since *z* + *e*^−*z*^ − 1 ≥ 0 for any *z* ≥ 0, we would have that *ϕ*^*′*^(*z*) ≤ 0. Since the function *ϕ*(*z*) is monotonically decreasing for *z* ≥ 0 we must have

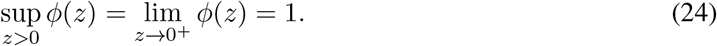

Observe that we can write *ϕ*^*′*^(*z*) as

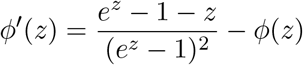

As the first term is always positive, we have *ϕ*^*′*^(*z*) ≥ −*ϕ*(*z*), which along with (24) shows that

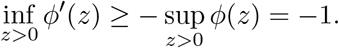

This completes the proof of the inequality (23) and concludes the proof of this lemma.

#### Proposition 1.

*For any u >* 0, *the integral*

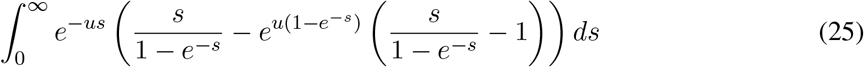

*has a positive value*.

*Proof*. Define a function

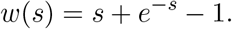

Note that the integral (25) can be rewritten as

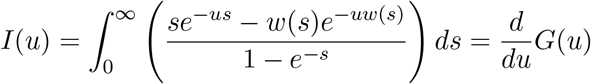

where

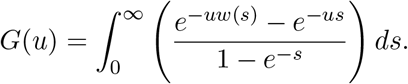

Hence in order to prove that *I*(*u*) is positive, we just need to prove that the function *G*(*u*) is monotonically increasing. This is what we show next.

Using the fact that

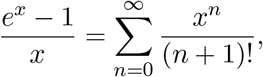

we obtain

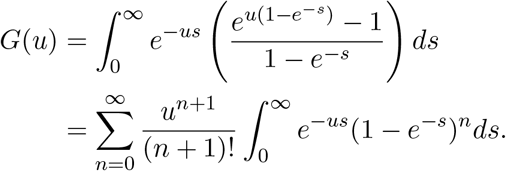

Applying the change of variable *t* = 1 − *e*^−*s*^, we get

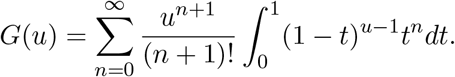

The integral on the right can be expressed in terms of the Gamma function Γ(*x*) as

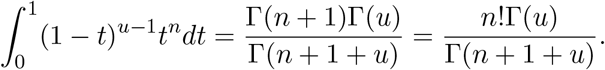

Substituting this we obtain

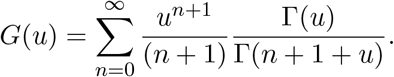

Since Γ(*x* + 1) = *x*Γ(*x*) we can express *G*(*u*) as

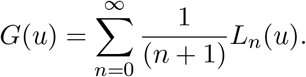

where

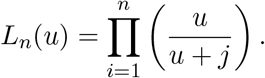

As each *L*_*n*_(*u*) is a product of positive monotonically increasing functions, the function *G*(*u*) is also monotonically increasing. This completes the proof of this proposition.

### 4. Proving monotonicity of the cDR

Finally we now prove the monotonicity of the cDR by proving (13). For this we shall use the integral expression (21).

Let us first deal with the case *u* ≤ *x*_0_. In this scenario (*u* − *x*_0_) ≤ 0 and since *e*^*s*−*T*^ ≤ 1 we have

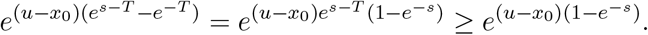

Therefore the integrand on the r.h.s. of (21) can be bounded below

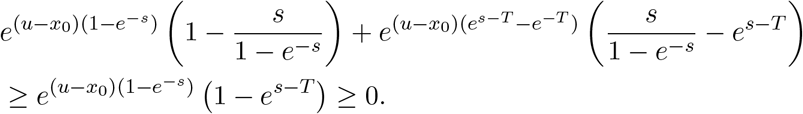

Hence the integrand in (21) is always non-negative which establishes the monotonicity of cDR for *u* ≤ *x*_0_.

We now consider the case *u > x*_0_. Note that in this case *x*_*u*_(*t*) (see (14)) is monotonically increasing from *x*_0_ to *u*. Hence *y*_*u*_(0) = 1 and

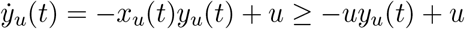

which allows us to conclude that *y*_*u*_(*t*) ≥ 1 for all *t* ≥ 0, by a simple comparison argument we now provide. Suppose that *x*_0_ ≤1. In this case we prove that for any positive *u*, the trajectory *t* ↦ ∂_*u*_*y*_*u*_(*t*) can only change its sign once, to go from positive to negative, and then it will stay negative and asymptotically approach 0. To see this define

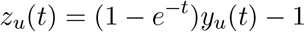

and then

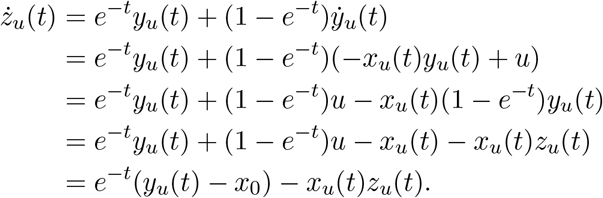

Since we have assumed *x*_0_ ≤ 1 we have the differential inequality

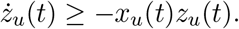

Hence, by the comparison argument, if there exists a *t*_1_ such that *z*_*u*_(*t*_1_) ≥ 0, then for all *t* ≥ *t*_1_ we have *z*_*u*_(*t*) ≥ 0 which also implies that

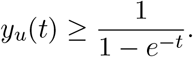

Now let *t*_1_ be the first zero of the trajectory ∂_*u*_*y*_*u*_(*t*). Observe that

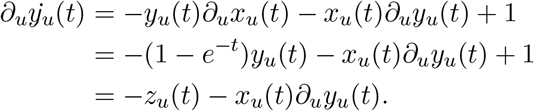

As *t*_1_ is the first zero of the trajectory ∂_*u*_*y*_*u*_(*t*) we have ∂_*u*_*y*_*u*_(*t*) *>* 0 for *t < t*_1_, and 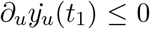 and ∂_*u*_*y*_*u*_(*t*_1_) = 0. Hence we must have *z*_*u*_(*t*_1_) ≥ 0 and by previous arguments *z*_*u*_(*t*) ≥ 0 for all *t* ≥ *t*_1_. Therefore in this interval *t* ∈ (*t*_1_, ∞) we have the inequality

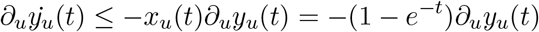

and since ∂_*u*_*y*_*u*_(*t*_1_) = 0, the comparison argument shows that ∂_*u*_*y*_*u*_(*t*) ≤ 0 for all *t* ≥ *t*_1_. Hence the trajectory *t* ↦ ∂_*u*_*y*_*u*_(*t*) can only change its sign once, to go from positive to negative. Therefore in order to prove 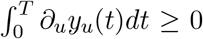 for any *T*, it suffices to prove that this holds in the limit *T* → ∞. Letting *T* → ∞ in (21) we arrive at

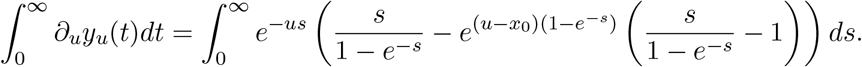

Since 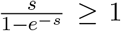 for all *s >* 0, in order to prove the positivity of this integral it suffices to prove its positivity for *x*_0_ = 0, i.e.

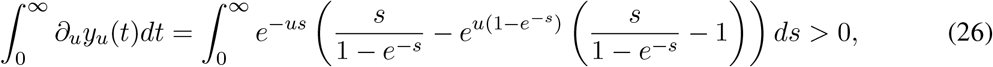

which holds due to Proposition 1.

We now come to the case where 1 *< x*_0_ *< u*. Set *κ* = (*u − x*_0_) and let *ρ*(*s*) and *f*_*κ*_(*s*) be the functions defined in the statement of Lemma 1. Note that (21) can be written in terms of this function as

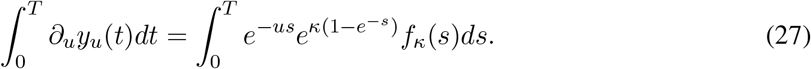

Moreover

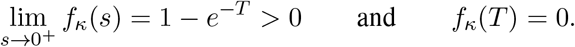

Lemma 1 proves that the function *f*_*κ*_(*s*) can only cross the *x*-axis at most once in the interval [0, *T*]. If it does cross then it goes from positive to negative and it stays negative till it becomes 0 at *s* = *T*. The same holds for the function 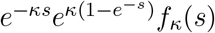. Hence if we can prove that

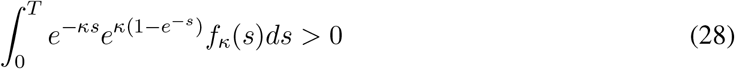

then, as κ < *u* it automatically implies that

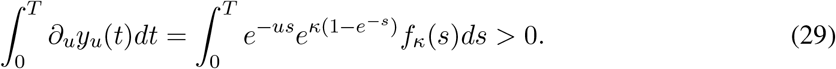

To see this let *s*\* be the time in [0, *T*] where the function *f*_*κ*_(*s*) crosses the *x*-axis. Then (28) implies that

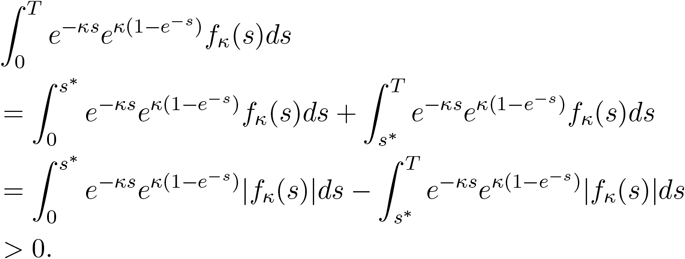

Therefore, using that *e*^−*us*^ ≥ *e*^−(*u*−*κ*)*s*^* *e*^−*κs*^ for *s* ≤ *s*\* and *e*^−*us*^ ≤ *e*^−(*u*−*κ*)*s*^* *e*^−*κs*^ for *s* ≥ *s*\*, we get

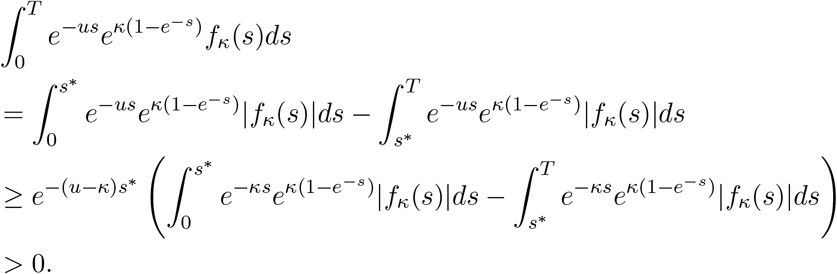

This shows that to prove the monotonicity result it suffices to prove (28), which is of course equivalent to proving the positivity of (21) for *u* = *κ* and *x*_0_ = 0. As mentioned above, this positivity follows from (26) which is shown in Proposition 1. This completes the proof of the cDR monotonicity result for IFFL1.

## 3 Unconditional monotonicity of both DR and cDR for IFFL2

Recall the description of IFFL2 from equations (7)-(8). Here we can easily prove that DR map *u* ↦ *y*_*u*_(*t*) is a monotonically increasing function of input *u*, and hence, the same holds for the cDR map. In order to prove this monotonicity it suffices to show that the partial derivative ∂_*u*_*y*_*u*_(*t*) is nonnegative for any *u* and *t*. Since *x*_*u*_(*t*) satisfies the linear ODE (7) we can solve for it explicitly to obtain

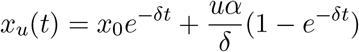

which also implies that

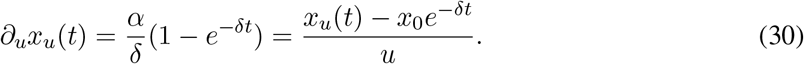

As *y*_*u*_(*t*) satisfies the ODE (8) we can differentiate it with respect to *u* to obtain an ODE for ∂_*u*_*y*_*u*_(*t*) as

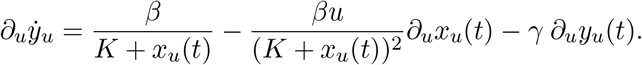

Substituting ∂_*u*_*x*_*u*_(*t*) from (30) and re-arranging we get

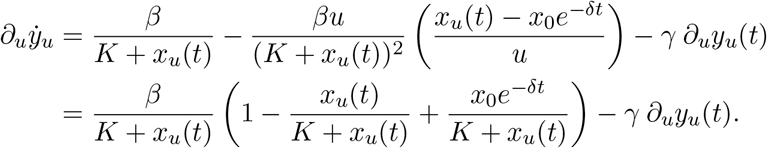

Since *K* ≥ 0, it is immediate that 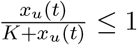, and so we have the differential inequality

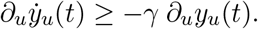

As ∂_*u*_*y*_*u*_(0) = 0, by the comparison argument it follows that ∂_*u*_*y*_*u*_(*t*) ≥ 0 for all *t* and *u*. This concludes the proof of the monotonicity result for IFFL2.

### 3.1 Monotone systems: an approach to show DR monotonicity of IFFL2

We may view the argument given for the monotonicity of the DR (and thus also the cDR) for IFFL2 in a more conceptual way, using the theory of monotone systems. Although the direct proof is simpler for this particular example, the more general approach is useful when analyzing other examples.

Monotone systems were introduced in the pioneering work of Smale, Smith, Hirsch, Mallet-Paret, Sell, and others [44, 45, 46, 47]. They have the property that larger initial conditions give rise to larger state trajectories, where “larger” is interpreted according to a specified order in the state variables. Special cases of monotone systems are obtained when the order is a coordinatewise order. For example in two dimensions, the “NorthEast” (NE) is defined by saying that a point (*x*_2_, *y*_2_) is larger than a point (*x*_1_, *y*_1_) if both *x*_2_ *> x*_1_ and *y*_2_ *> y*_1_, that is, if it is to the “north” and “east” (higher, to the right) of the second point; similarly the “NorthWest” order would be defined by asking that *x*_2_ *< x*_1_ and *y*_2_ *> y*_1_. (Note that these are “partial orders” in the sense that two vectors may not be comparable: for example neither (0, 1) nor (1, 0) is larger than the other in the NE order.) The generalization to external inputs and outputs [48] enabled the development of a network interconnection theory as well as leading to conclusions regarding the effect of inputs: for example, monotonic inputs result in monotonic transient behavior [49]. This means that the DR (and thus also the cDR) will always be monotonic, for monotone systems.

These developments led to monotone systems playing a key role in analyzing the global behavior of dynamical systems in various areas of engineering as well as biology [50]. What makes monotone system theory so useful is that there are ways to check monotonicity without solving a set of differential equations 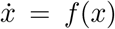. For example, for the *n*-dimensional analog of the NE order one requires that the off-diagonal terms of the Jacobian matrix of *f* should all be nonnegative, and a similar condition holds if there are inputs. More generally, monotonicity with respect to some (not necessarily the NE) coordinatewise order requires that all loops in the interaction graph obtained from the off-diagonal terms of the Jacobian should have a net positive sign. See e.g. [51] for more discussion and examples.

When trying to apply monotone systems theory to our three paradigmatic examples IFFL1, IFFL2, and NFL, an obvious problem arises: these systems are not monotone with respect to any possible coordinatewise order, as incoherent feedforward loops and negative feedback loops contradict the positive loop condition. However, in the case of the IFFL2 system, which we reproduce here for convenience:

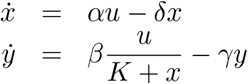

there is a Lie group of symmetries or “equivariances” that preserve the output. These equivariances were the main object of study in the work in [8] on scale invariance, and where key to the analysis of an immunology model in [29]. Specifically, when *K* = 0 the discussion in Section 1.6 showed that the equations do not change under the one-parameter Lie group of transformations (*u, x, y*) ↦ (*pu, px, y*), and in particular scaling *u* and *x* by the same constant does not alter the dynamics of *y*. This suggests introducing the new variable *p* := *u/x*. Using the variables *p* and *y*, the equations become:

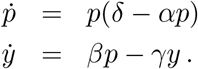

This is a monotone system, because the only off-diagonal term in the Jacobian is *β >* 0. Therefore the trajectories depend monotonically on *p*(0) = *u/x*_0_, and hence also depend monotonically on *u*. A similar argument works for *K >* 0, but now the FCD property fails and the equivariance will provide merely an embedding into a monotone system rather than an equivalence. Indeed, let *p* := *u/*(*K* + *x*). Now the *p* equation is no longer decoupled from *x*. However, we can look at the following extended system (we add an equation 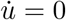 to convert the external input into a state variable):

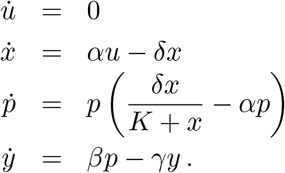

The off-diagonal elements of the Jacobian are *α, β*, and *Kδp/*(*K* + *x*)^2^, all positive. Thus the extended system is monotone, and therefore all variables, and in particular *y*, depend monotonically on *u*. This shows monotonicity of the DR, as claimed.

## 4 Conditional monotonicity of the cDR for IFB

Recall the description of IFB from equations (9)-(10). We shall assume that *K* = 0 for convenience. We scale time by *γ*^−1^ and the state values by a ratio to define

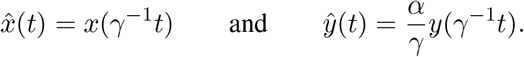

Then we can write this system as

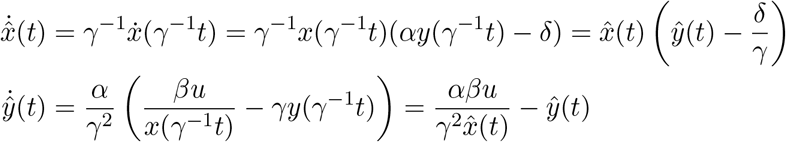

Therefore if we define

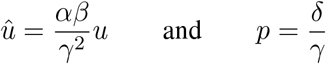

then 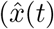, *ŷ*(*t*)) satisfies the ODEs

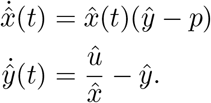

with initial values 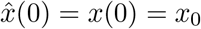 and 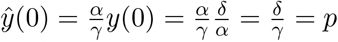. Henceforth we shall drop the hats for notational convenience and suppose that the dynamics is given by

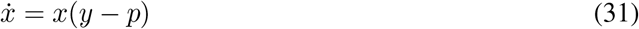

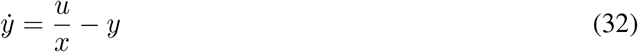

with initial values *x*(0) = *x*_0_ and *y*(0) = *p*.

Note that *p* is both the initial state and the steady-state value of the output variable *y*_*u*_(*t*). We shall reformulate the dynamics in terms of the “error”

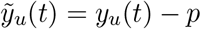

and variable *z*_*u*_(*t*) defined as

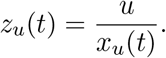

Then dynamics of *z*_*u*_(*t*) is given by

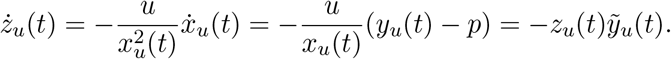

Hence we can solve for *z*_*u*_(*t*) as

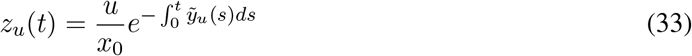

and the dynamics of the error is given by

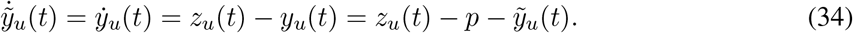

Henceforth, instead of examining the monotonicity of the original cDR map, we shall equivalently examine the monotonicity of the error cDR map given by

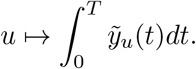

### 1. Steady-state analysis

We proved earlier that the steady state is stable. Thus 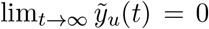 and so lim_*t*→∞_ *z*_*u*_(*t*) = *p*. Therefore expression (33) implies that

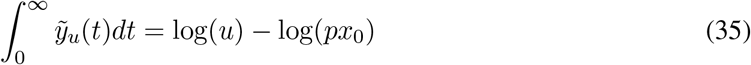

which is a monotonically increasing function of *u*. In particular, by differentiating with respect to *u* we obtain

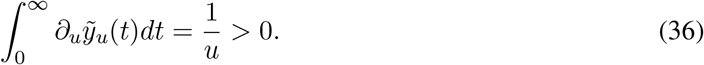

If this positivity holds for any finite time-interval then we shall have the monotonicity of the error cDR map. We shall show that this monotonicity does not always hold and identify a sufficient condition under which it does. For this we shall connect this problem to a harmonic oscillator with a time-varying frequency.

### 2. Connection to a harmonic oscillator

Differentiating the error equation (34) with respect to *t* and using that 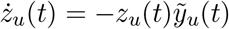 we get

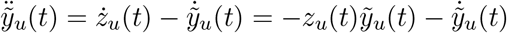

which means that the error 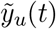 satisfies the equation for a ‘damped’ harmonic oscillator with a non-constant frequency

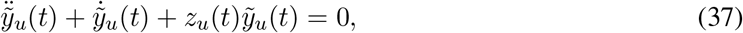

with initial conditions 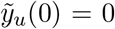 and 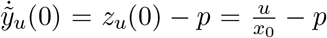. Let us define the ‘frequency’ and Hamiltonian for this oscillator as

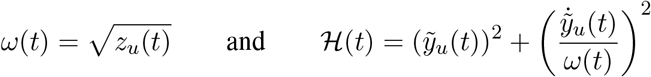

respectively. Then their dynamics can be derived as

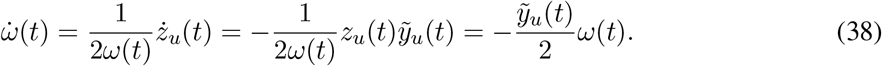

Using (37) we obtain

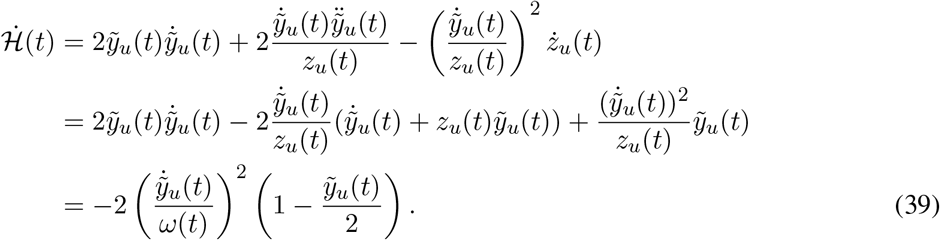

Observe that the error 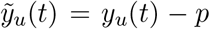 in our adapting circuit goes to 0 as *t* → ∞. Equation (39) shows that when this error is below 2, the Hamiltonian is decreasing. Note that the initial value of this Hamiltonian is

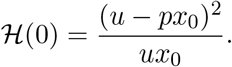

This brings us to a proposition that shows that if this value is less than 4, then the error 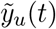 remains below 2 at all times.

#### Proposition 2.

*Suppose that the following holds*

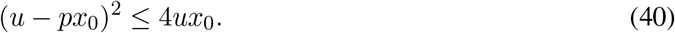

*Then we must have that* 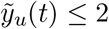 *for all t* ≥ 0.

*Proof*. Condition (40) implies that ℋ (0) ≤ 4. Recall that 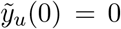. Let *t*_1_ be the first time 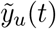 reaches 2, i.e. 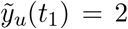 and 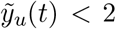 for all *t < t*_1_. Then due to (39) the Hamiltonian must be decreasing in the interval [0, *t*_1_], and so ℋ (*t*_1_) *<* ℋ (0) ≤ 4. But this is a contradiction since 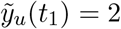 and so by definition ℋ (*t*_1_) ≥ 4. Hence *t*_1_ = ∞ which means that 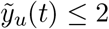 for all *t* ≥ 0.

### 3. Conditional monotonicity result

Now let us substitute *z*_*u*_(*t*) from (33) into eqn. (34) to obtain

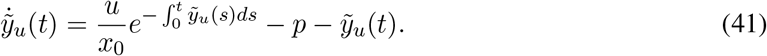

Differentiating this with respect to *u* we get

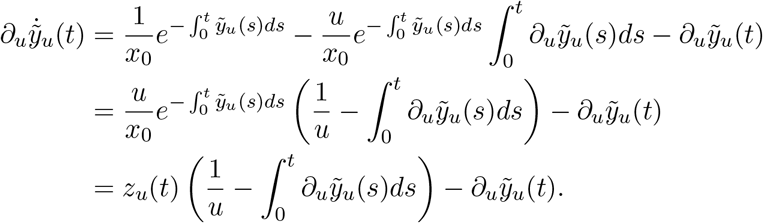

This shows that if we define *β*(*t*) as

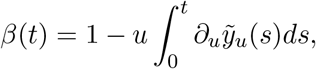

then *β*(*t*) also satisfies the following second-order ODE for the damped harmonic oscillator (37) with initial conditions *β*(0) = 1 and 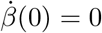. To prove the monotonicity we need to show that

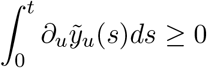

which is equivalent to proving that

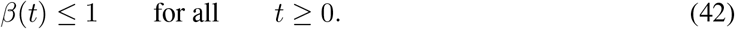

Again we can definite the Hamiltonian as

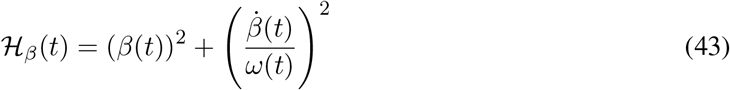

and it will have the same dynamics as before

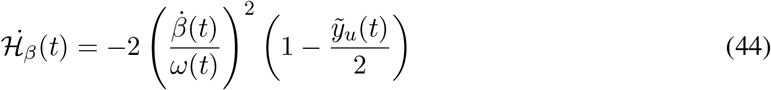

which we can also write as

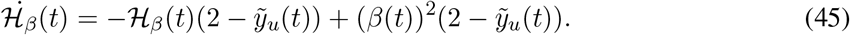

We now come to our main result for the IFB example, which proves monotonicity of the error cDR under condition (40).

#### Proposition 3.

*Suppose that condition* (40) *holds. Then the map*

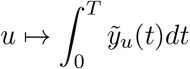

*is monotonically increasing for any T >* 0.

*Proof*. When (40) holds, we have that the error 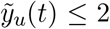 for all *t* ≥ 0 due to Proposition 2. This fact along with (44) implies that ℋ_*β*_(*t*) ≤ ℋ_*β*_(0) = 1 for *t* ≥ 0. Therefore (42) holds which proves the required monotonicity property.

### 4. Numerical illustration of non-monotonicity

When condition (40) does not hold, we do not have any analytical approach for checking the monotonicity property. Therefore we need to rely on numerical simulations. For any given *u*, the monotonicity condition (42) can fail at any *t* ≥ 0. However as we cannot check this condition for all *t* ≥ 0, we first show that if this monotonicity fails then it has to fail in the finite time-interval [0, *T*_*u*_] where *T*_*u*_ is defined by

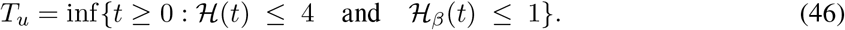

Note that *T*_*u*_ is finite because both ℋ(*t*) and ℋ_*β*_(*t*) are nonnegative quantities converging to zero as *t* → ∞. This is due to stability and the fact that (36) implies that *β*(*t*) → 0 as *t* → ∞.

#### Proposition 4.

*Let T*_*u*_ *be the finite time defined by* (46). *Then we must have β*(*t*) ≤ 1 *for all t* ≥ *T*_*u*_ *which is the same as saying that u* 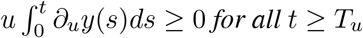.

*Proof*. Pick a small *ϵ >* 0 and define

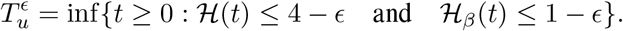

Note that for the same reason that *T*_*u*_ is finite, 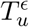 must also be finite. Moreover as *ϵ* decreases the set *{t* ≥ 0 : ℋ (*t*) ≤ 4 − *ϵ* and ℋ_*β*_(*t*) ≤ 1 − *ϵ}* gets bigger and consequently its infimum, which is 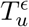, gets smaller. Therefore 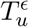 decreases monotonically as *ϵ* decreases and since ℋ (*t*) and ℋ_*β*_(*t*) are continuous functions of time we must have

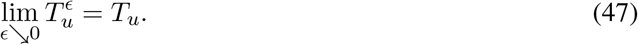

By the definition of 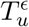 we have that 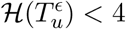 which implies that 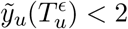. Let 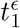 be the first time after 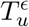 such that 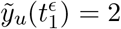 and consequently 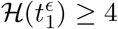. On the open interval 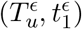, ℋ(*t*) must be decreasing due to (39) and so we should have 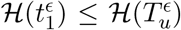. However this leads to a contradiction because 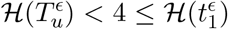. Therefore 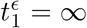 which means that 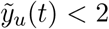 for all 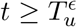. This also means that ℋ_*β*_(*t*) in decreasing in the interval 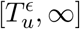 due to (44). Since 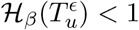 this implies that *β*(*t*) *<* 1 for all 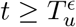. Using the continuity of *β* and the limit (47) we can conclude that *β*(*t*) ≤ 1 for all *t* ≥ *T*_*u*_. This completes the proof of this proposition.

Figure 8 illustrates this proposition for three values of *u*. One can observe that 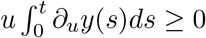 holds for all *t* ≥ *T*_*u*_ for each *u*-value.

**Figure 8:**
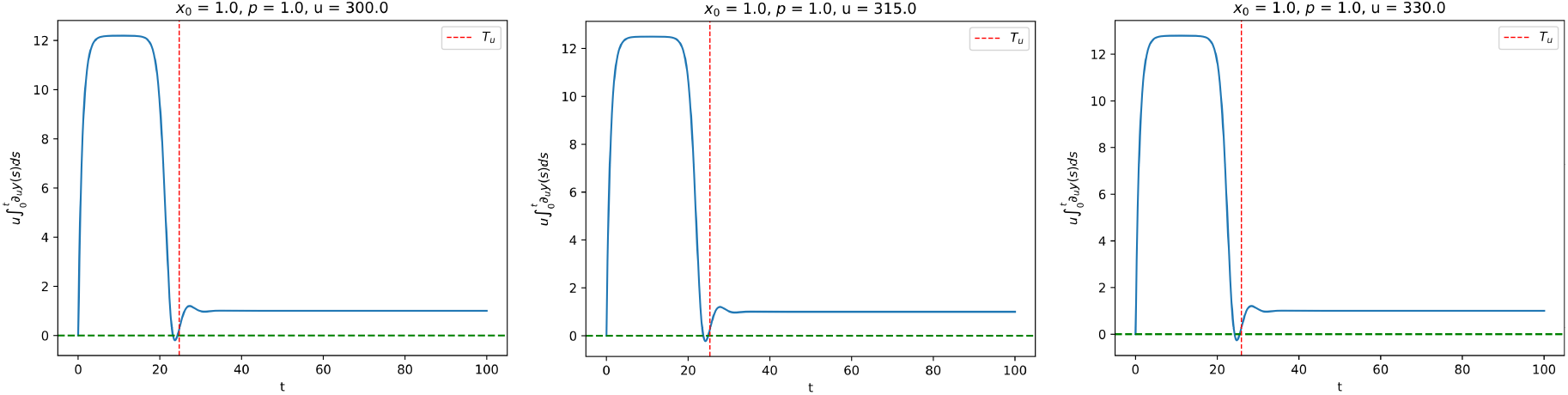
Plot of the map 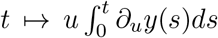 for three values of *u*. Note that for each *u*, we have 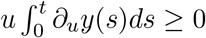 for all *t* ≥ *T*_*u*_.

We can now numerically solve the system (*z*_*u*_(*t*), *y*_*u*_(*t*), *β*(*t*)) in the interval [0, *T*_*u*_] and check if *β*(*t*) exceeds 1 or not. For any *t* such that *β*(*t*) *>* 1 we would have 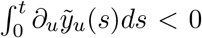 which would imply non-monotonicity. For any given *u* we define the non-monotonicity score as

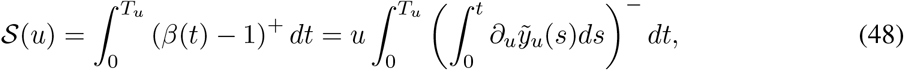

where *a*^+^ (resp. *a*^−^) denotes the positive (resp. negative) part of *a*. In Figure 9 we plot the non-monotonicity score *𝒮*(*u*) and the time *T*_*u*_ for a range of *u*-values for *x*_0_ = *p* = 1. Notice that *T*_*u*_ is monotonically increasing with *u*, while the non-monotonicity score 𝒮 (*u*) stays at zero till *u* ≈ 220 and then it gradually starts increasing. This shows that for higher values of *u* the map

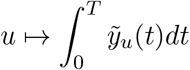

is always non-monotonic for some *T* which will increase as *u* increases. Figure 10 illustrates this non-monotonicity for some values of *p* and *T*.

**Figure 9:**
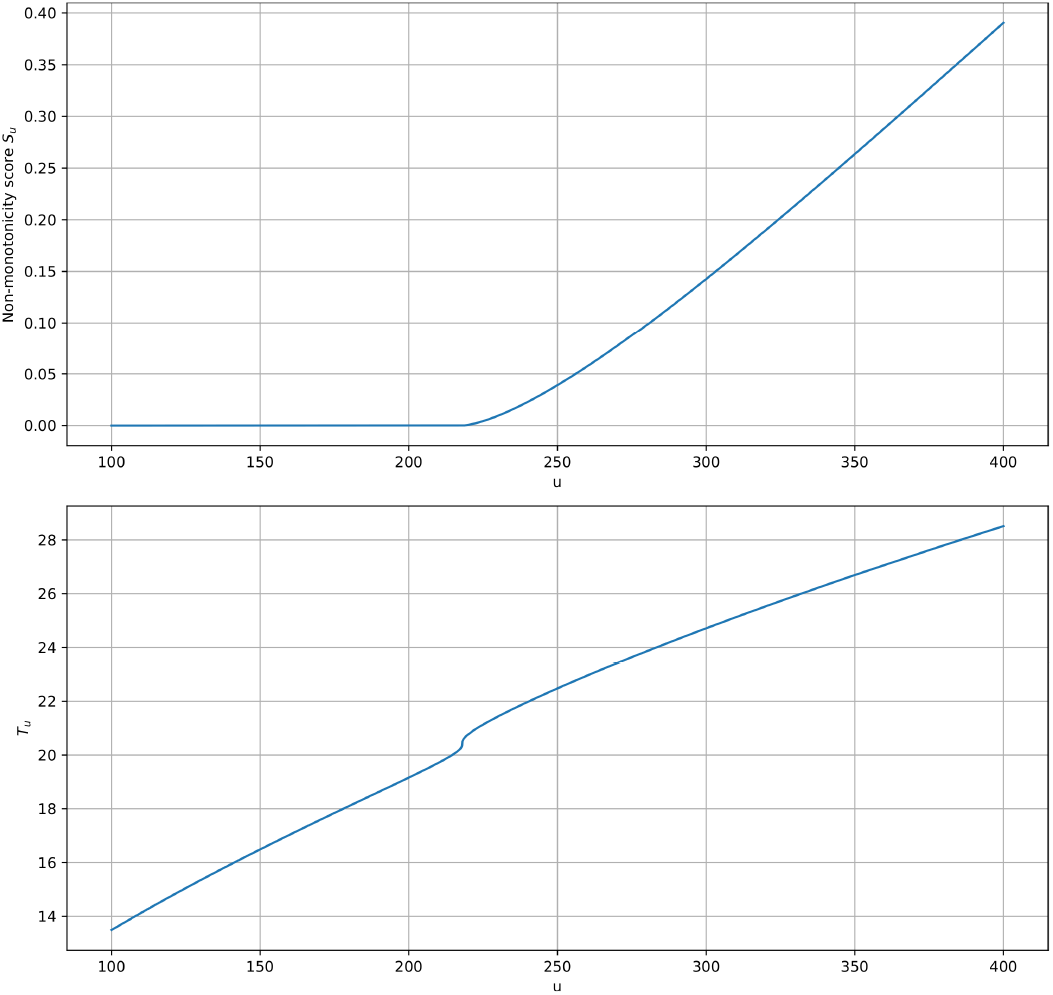
Plot of the maps *u* ↦ 𝒮(*u*) and *u* ↦ *T* (*u*) with *x*_0_ = *p* = 1.0.

**Figure 10:**
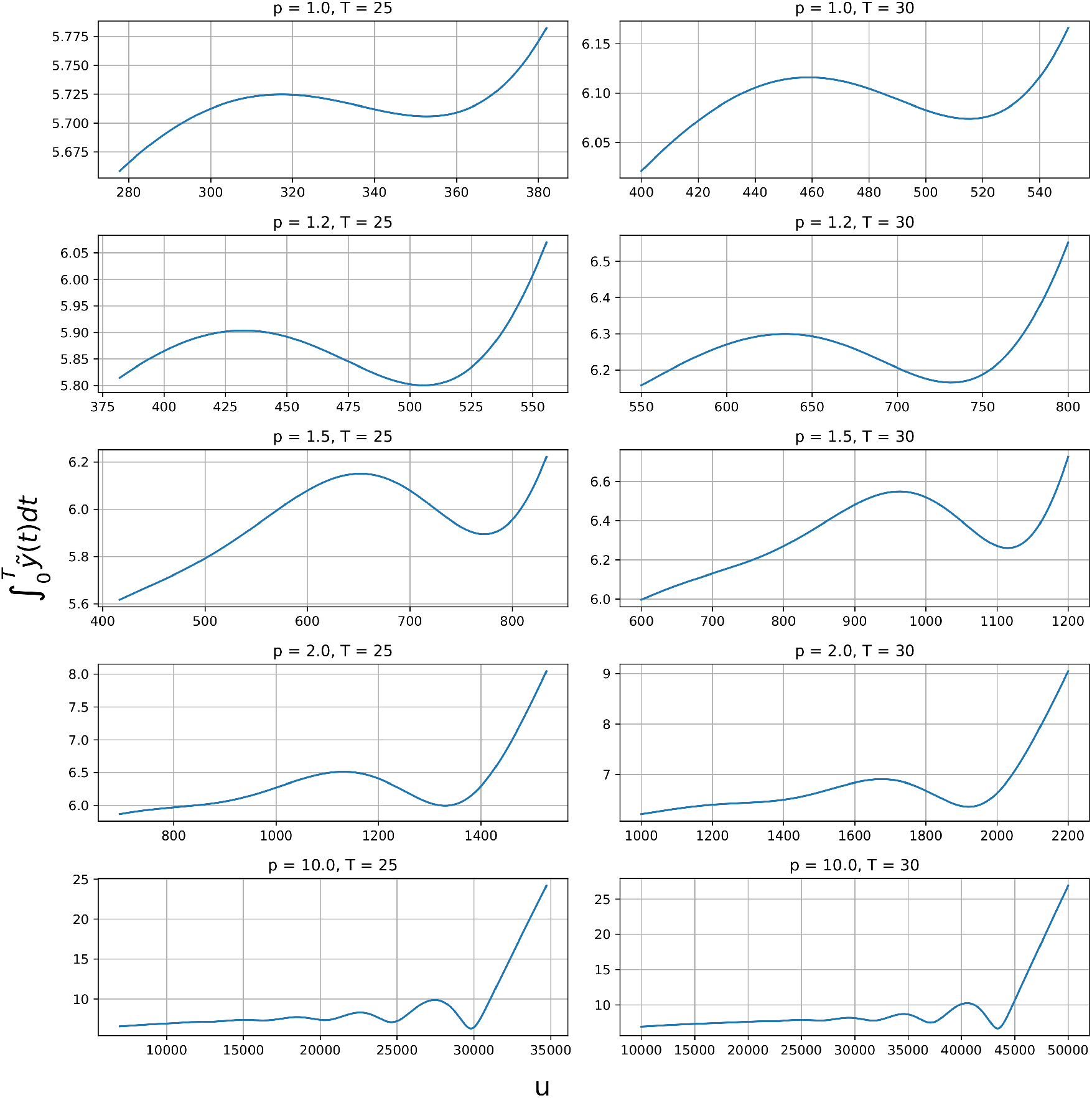
Plot of the map 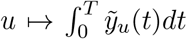 for some values of *p* and *T*. Note that this map exhibits non-monotonicity in these cases.

## 5 Conclusions

We introduced the notion of cDR, and went on to show mathematically that both the incoherent feedback loop IFFL1 and IFFL2 motifs can only produce monotonic cumulative dose responses, even if (for IFFL1) the dose response itself may be non-monotonic. On the other hand, we also established conditions under which the integral feedback mechanism IFB can produce a non-monotonic cDR.

One may view our study of the cDR work as an addition to the toolkit of mathematical methods for model discrimination and invalidation, in a spirit similar to the work in [52] that infers the existence of IFFLs or negative feedback loops when time responses are non-monotonic, or to the work in [10] on using periodic inputs in order to rule out IFFLs as the basis of adaptation.

## 6 Acknowledgments

The authors thank Dr. Omer Dushek, from the University of Oxford, for generously sharing experimental data. EDS’s research was supported in part by grant AFOSR FA9550-21-1-0289. AG’s research was supported in part by the Swiss National Science Foundation (SNSF) grant no. 216505.

## References

[1] N. Barkai and S. Leibler.Robustness in simple biochemical networks. Nature, 387(6636):913– 917, Jun 1997.

[2] Graeme W. Davis. Homeostatic control of neural activity: From phenomenology to molecular design. Annual Review of Neuroscience, 29(Volume 29, 2006):307–323, 2006.

[3] B. Wark, B. N. Lundstrom, and A. Fairhall. Sensory adaptation. Current Opinion in Neurobiology, 17(4):423–429, 2007.

[4] D. Muzzey, C.A. Gòmez-Uribe, J. T. Mettetal, and A. van Oudenaarden. A systems-level analysis of perfect adaptation in yeast osmoregulation. Cell, 138(1):160–171, Jul 2009.

[5] M.H. Khammash. Perfect adaptation in biology. Cell Systems, 12(6):509–521, Jun 2021.

[6] N. Trendel, P. Kruger, S. Gaglione, J. Nguyen, J. Pettmann, E.D. Sontag, and O. Dushek. Perfect adaptation of CD8+ T cell responses to constant antigen input over a wide range of affinity is overcome by costimulation. Science Signaling, 14:eaay9363, 2021.

[7] O. Shoval, L. Goentoro, Y. Hart, A. Mayo, E.D. Sontag, and U. Alon. Fold change detection and scalar symmetry of sensory input fields. Proc Natl Acad Sci USA, 107:15995–16000, 2010.

[8] O. Shoval, U. Alon, and E.D. Sontag. Symmetry invariance for adapting biological systems. SIAM Journal on Applied Dynamical Systems, 10:857–886, 2011.

[9] W. Ma, A. Trusina, H. El-Samad, W. A. Lim, and C. Tang. Defining network topologies that can achieve biochemical adaptation. Cell, 138(4):760–773, 2009.

[10] S. J. Rahi, J. Larsch, K. Pecani, N. Mansouri, A. Y. Katsov, K. Tsaneva-Atanasova, E. D. Sontag, and F. R. Cross. Oscillatory stimuli differentiate adapting circuit topologies. Nature Methods, 14:1010–1016, 2017.

[11] P. Yu and E.D. Sontag. A necessary condition for non-monotonic dose response, with an application to a kinetic proofreading model. In Proc. 2024 63rd IEEE Conference on Decision and Control (CDC), 2024. To appear. Note: Extended version in arXiv.

[12] J.J. Tyson, K. Chen, and B. Novak. Sniffers, buzzers, toggles, and blinkers: dynamics of regulatory and signaling pathways in the cell. Curr. Opin. Cell. Biol., 15:221–231, 2003.

[13] E.D. Sontag. Remarks on feedforward circuits, adaptation, and pulse memory. IET Systems Biology, 4:39–51, 2010.

[14] U. Alon. An Introduction to Systems Biology: Design Principles of Biological Circuits. Chapman & Hall, 2006.

[15] S. Semsey, S. Krishna, K. Sneppen, and S. Adhya. Signal integration in the galactose network of Escherichia coli. Mol. Microbiol., 65:465–476, Jul 2007.

[16] F.-D. Xu, Z.-R. Liu, Z.-Y. Zhang, and J.-W. Shen. Robust and adaptive microRNA-mediated incoherent feedforward motifs. Chinese Physics Letters, 26(2):028701–3, February 2009.

[17] S. Sasagawa, Y. Ozaki, K. Fujita, and S. Kuroda. Prediction and validation of the distinct dynamics of transient and sustained ERK activation. Nat. Cell Biol., 7:365–373, Apr 2005.

[18] T. Nagashima, H. Shimodaira, K. Ide, T. Nakakuki, Y. Tani, K. Takahashi, N. Yumoto, and M. Hatakeyama. Quantitative transcriptional control of ErbB receptor signaling undergoes graded to biphasic response for cell differentiation. J. Biol. Chem., 282:4045–4056, Feb 2007.

[19] P. Menè, G. Pugliese, F. Pricci, U. Di Mario, G. A. Cinotti, and F. Pugliese. High glucose level inhibits capacitative Ca2+ influx in cultured rat mesangial cells by a protein kinase C-dependent mechanism. Diabetologia, 40:521–527, May 1997.

[20] R. Nesher and E. Cerasi. Modeling phasic insulin release: immediate and time-dependent effects of glucose. Diabetes, 51 Suppl 1:S53–59, Feb 2002.

[21] M. P. Mahaut-Smith, S. J. Ennion, M. G. Rolf, and R. J. Evans. ADP is not an agonist at P2X(1) receptors: evidence for separate receptors stimulated by ATP and ADP on human platelets. Br. J. Pharmacol., 131:108–114, Sep 2000.

[22] S. Marsigliante, M. G. Elia, B. Di Jeso, S. Greco, A. Muscella, and C. Storelli. Increase of [Ca(2+)](i) via activation of ATP receptors in PC-Cl3 rat thyroid cell line. Cell. Signal., 14:61–67, Jan 2002.

[23] L. Yang and P.A. Iglesias. Positive feedback may cause the biphasic response observed in the chemoattractant-induced response of dictyostelium cells. Systems Control Lett., 55(4):329–337, 2006.

[24] A. Levchenko and P.A. Iglesias. Models of eukaryotic gradient sensing: Application to chemotaxis of amoebae and neutrophils. Biophys J., 82:50–63, 2002.

[25] L. A. Ridnour, A. N. Windhausen, J. S. Isenberg, N. Yeung, D. D. Thomas, M. P. Vitek, D. D. Roberts, and D. A. Wink. Nitric oxide regulates matrix metalloproteinase-9 activity by guanylyl-cyclase-dependent and -independent pathways. Proc. Natl. Acad. Sci. U.S.A., 104:16898–16903, Oct 2007.

[26] J. Tsang, J. Zhu, and A. van Oudenaarden. MicroRNA-mediated feedback and feedforward loops are recurrent network motifs in mammals. Mol. Cell, 26:753–767, Jun 2007.

[27] A. Kremling, K. Bettenbrock, and E. D. Gilles. A feed-forward loop guarantees robust behavior in escherichia coli carbohydrate uptake. Bioinformatics, 24:704–710, 2008.

[28] E. Voit, A. R. Neves, and H. Santos. The intricate side of systems biology. Proc. Natl. Acad. Sci. U.S.A., 103:9452–9457, Jun 2006.

[29] E.D. Sontag. A dynamical model of immune responses to antigen presentation predicts different regions of tumor or pathogen elimination. Cell Systems, 4:231–241, 2017.

[30] L. Bleris, Z. Xie, D. Glass, A. Adadey, E.D. Sontag, and Y. Benenson. Synthetic incoherent feed-forward circuits show adaptation to the amount of their genetic template. Molecular Systems Biology, 7:519–, 2011.

[31] T.H. Segall-Shapiro, E. D. Sontag, and C. A. Voigt. Engineered promoters enable constant gene expression at any copy number in bacteria. Nature Biotechnology, 36:352–358, 2018.

[32] D. Kim, Y. K. Kwon, and K. H. Cho. The biphasic behavior of incoherent feed-forward loops in biomolecular regulatory networks. Bioessays, 30:1204–1211, Nov 2008.

[33] T. M. Yi, Y. Huang, M. I. Simon, and J. Doyle. Robust perfect adaptation in bacterial chemotaxis through integral feedback control. Proc. Natl. Acad. Sci. U.S.A., 97:4649–4653, 2000.

[34] K. V. Venkatesh, S. Bhartiya, and A. Ruhela. Mulitple feedback loops are key to a robust dynamic performance of tryptophan regulation in Escherichia coli. FEBS Letters, 563:234–240, 2004.

[35] H. El-Samad, J. P. Goff, and M. Khammash. Calcium homeostasis and parturient hypocalcemia: An integral feedback perspective. J. Theor. Biol., 214:17–29, 2002.

[36] P. Miller and X. J. Wang. Inhibitory control by an integral feedback signal in prefrontal cortex: A model of discrimination between sequential stimuli. Proc. Natl. Acad. Sci., 103:201–206, 2006.

[37] D.K. Agrawal, R. Marshall, V. Noireaux, and E.D. Sontag. In vitro implementation of robust gene regulation in a synthetic biomolecular integral controller. Nature Communications, 10:1–12, 2019.

[38] S. K. Aoki, G. Lillacci, A. Gupta, A. Baumschlager, D. Schweingruber, and M. Khammash. A universal biomolecular integral feedback controller for robust perfect adaptation. Nature, 570(7762):533–537, Jun 2019.

[39] C. Briat, A. Gupta, and M. Khammash. Antithetic integral feedback ensures robust perfect adaptation in noisy biomolecular networks. Cell systems, 2(1):15–26, 2016.

[40] M. Bin, J. Huang, A. Isidori, L. Marconi, M. Mischiati, and E. D. Sontag. Internal models in control, bioengineering, and neuroscience. Annual Review of Control, Robotics, and Autonomous Systems, 5:20.1–20.25, 2022.

[41] A. Gupta and M. Khammash. Universal structural requirements for maximal robust perfect adaptation in biomolecular networks. Proceedings of the National Academy of Sciences, 119(43):e2207802119, 2022.

[42] Y. Hirono, A. Gupta, and M. Khammash. Complete characterization of robust perfect adaptation in biochemical reaction networks. arXiv preprint 2307.07444, 2023.

[43] E.D. Sontag. Mathematical Control Theory. Deterministic Finite-Dimensional Systems, volume 6 of Texts in Applied Mathematics. Springer-Verlag, New York, second edition, 1998.

[44] S. Smale. On the differential equations of species in competition. J. Math. Bio, 3:5–7, 1976.

[45] M.W. Hirsch. Stability and convergence in strongly monotone dynamical systems. Reine und Angew. Math, 383:1–53, 1988.

[46] H. Smith. Monotone Dynamical Systems: An Introduction to the Theory of Competitive and Cooperative Systems, Mathematical Surveys and Monographs, vol. 41. AMS, Providence, RI, 1995.

[47] J. Mallet-Paret and G. Sell. The Poincare-Bendixson theorem for monotone cyclic feedback systems with delay. Journal of Differential Equations, 125:441–489, 1996.

[48] D. Angeli and E.D. Sontag. Monotone control systems. IEEE Trans. Automat. Control, 48(10):1684–1698, 2003.

[49] D. Angeli and E.D. Sontag. Behavior of responses of monotone and sign-definite systems. In K. Hüper and Jochen Trumpf, editors, Mathematical System Theory - Festschrift in Honor of Uwe Helmke on the Occasion of his Sixtieth Birthday, pages 51–64. CreateSpace, 2013.

[50] D. Angeli, J. E. Ferrell, and E.D. Sontag. Detection of multistability, bifurcations, and hysteresis in a large class of biological positive-feedback systems. Proc Natl Acad Sci USA, 101(7):1822–1827, 2004.

[51] E.D. Sontag. Monotone and near-monotone biochemical networks. Systems and Synthetic Biology, 1:59–87, 2007.

[52] J.A. Ascensao, P. Datta, B. Hancioglu, E.D. Sontag, M.L. Gennaro, and O.A. Igoshin. Non-monotonic response dynamics of glyoxylate shunt genes in mycobacterium tuberculosis. PLoS Computational Biology, 12:e1004741, 2016.

